# Home is Where the Heterogeneity Is: Housing Facility-level Differences in the Gut Microbiome and Metabolic Phenotype Confound Arsenic Effects on Glucose Homeostasis in Male Mice

**DOI:** 10.64898/2026.08.03.742222

**Authors:** Judy Malas, Lidan Zhao, Michael Landeche, Ashley M. Sidebottom, Jessica Little, Jarrad Hampton-Marcell, Robert M. Sargis

## Abstract

Inorganic arsenic (iAs) exposure is linked to impaired glucose homeostasis and type 2 diabetes, yet the magnitude and direction of reported effects vary substantially across studies and populations. The gut microbiome is both a target and a mediator of arsenic toxicity, suggesting that pre-exposure community composition may modulate the development of metabolic dysfunction. To test this, we conducted parallel 50 ppm iAs drinking-water exposures in male C57BL/6J mice at two animal facilities. Results were compared across facilities for metabolic phenotypes, hepatic arsenic levels, targeted and untargeted metabolomics, and shotgun metagenomics. Hepatic arsenic confirmed comparable exposure at both sites; however, the housing facility explained more variance than the iAs treatment group across every data layer. Baseline microbial communities and metabolic phenotypes at each institution differed, and this difference propagated into the iAs treatment effect. Critically, iAs exposure impaired glucose clearance at one site while trending toward improvement at the other. Facility explained 19 to 26% of variance in microbiome, bile acid, polar, and untargeted metabolite ordinations, while iAs treatment did not reach significance. A random forest classifier identified the facility with 96% cross-validated accuracy from 22 microbial species, whereas treatment classification did not exceed 67% accuracy. Functional metagenomic analyses revealed nearly 11,733 (63%) of genes were differentially abundant between facilities compared 139 with iAs treatment. Our results indicate that identical genetics and exposure may produce differential metabolic outcomes on different microbial backgrounds. Characterizing the baseline microbiome and metabolome is therefore critical both for identifying which individuals are most susceptible to the metabolic effects of arsenic exposure and for potentially reducing the risk of exposure through modulation of the gut microbiome.

## Introduction

Growing evidence suggests that exposure to endocrine-disrupting chemicals such as non-essential metals, persistent organic pollutants, organophosphates, and phthalates is a substantial contributor to adverse health effects (Kirkley and Sargis 2014; La Merrill et al. 2024; Münzel et al. 2023; Shrivastav, Swetanshu, and Singh 2024; Velmurugan et al. 2017). The gut microbiome is a central factor in mediating host health and is a key intermediary linking toxicant exposures to health outcomes (Ahn and Hayes 2020; Jorgensen et al. 2025). Therefore, numerous experimental studies on toxicant exposure have included gut microbiome and metabolomic measurements to understand the role of the microbiome in toxicant response.

Exposure to inorganic arsenic (iAs) has been the focus of several epidemiologic and experimental animal model studies. Globally, up to 220 million people are exposed to inorganic arsenic (iAs) from contaminated drinking water (Podgorski and Berg 2020). Arsenic is a class 1 carcinogen that is also linked to several chronic metabolic diseases, including prediabetes, gestational diabetes mellitus, and type 2 diabetes mellitus in epidemiological studies (Kim et al. 2022; Kuo et al. 2013; Mimoto, Nadal, and Sargis 2017; Salmeri et al. 2020; Spaur et al. 2024; Weiss, Sun, Jackson, Turyk, Wang, Brown, Aguilar, Brown, et al. 2024; Wu et al. 2024). Moreover, animal models and cell-based models confirm the capacity of iAs to disrupt glucose homeostasis (Kirkley et al. 2018; Li et al. 2020; Paul et al. 2011).

Importantly, rodent studies also indicate that iAs exposure alters gut microbiome composition and function, modifies metabolite profiles, and induces gut barrier dysfunction (Chi et al. 2017; Chiocchetti et al. 2019; A. Domene et al. 2023; Adrián Domene et al. 2023; Dong, Luo, and Zhang 2024; Li et al. 2021; Lu et al. 2014). Furthermore, iAs-induced phenotypic changes are transferable following fecal microbiome transplant, as mice gavaged with stool from arsenic-exposed mice exhibit severe inflammatory cell infiltration in liver tissues, implicating disruptions in the gut microbiome as a mediator of arsenic toxicity (Dong et al. 2024). However, results differ in the compositional alterations induced by arsenic exposure, and large within-treatment microbiome variability has been observed (Richardson et al. 2018). In addition, while arsenic-induced dysbiosis can negatively impact host health, the gut microbiome can also protect the host by transforming the molecular speciation of arsenic and facilitating its excretion (Coryell et al. 2018). Together, these observations suggest that the composition of the gut microbiome is a key determinant of how the host responds to iAs exposure.

While environmental toxicant exposures can alter the microbiome, several other factors, including genotype, lifestyle, diet, stress, and exposure to external microbiota, play a major role in shaping the gut microbiome (Grieneisen et al. 2021; Rothschild et al. 2018). Therefore, the specific starting composition of the gut microbiome likely influences the toxicological response to environmental perturbations. In this work, we conducted two parallel iAs exposure treatment experiments at two different animal facilities and examined the role of the gut microbiome in modifying toxicant-associated metabolic outcomes.

## Methods

### Animal care and arsenic exposure

Seven-week-old male C57BL/6J mice (Jackson Laboratory, Bar Harbor, Maine) were group-housed in facilities either at Institution A (InstA) or Institution B (InstB). Mice in both facilities had *ad libitum* access to standard chow (Teklad 2918) and water. Per facility, mice were grouped into iAs exposure or control groups. Mice in the iAs treatment groups were provided bottles of water containing 50 ppm sodium arsenite (As^3+^) for drinking. At InstA, 10 mice per treatment group (n = 20) were housed under a 12-hour light-dark cycle, and the source water was reverse-osmosis-purified. At InstB, eight mice per treatment group (n = 16) were housed under 14-hour light, 10-hour dark cycles, and given autoclaved tap water. Both institutions were in the same city and received municipal water from the same source.

### Fasting glucose, fasting insulin, and glucose tolerance tests

Glucose tolerance tests (GTT) were performed at 8 weeks and 12 weeks of age. Mice were fasted for 6 h and administered 2 g/kg glucose by I.P. injection. Blood glucose from the tail vein was measured at 0, 10, 20, 30, 60, 90, and 120 min using a FreeStyle Freedom Lite meter and test strips (Abbott, Alameda, CA, USA). An aliquot of 0 min blood was collected into a heparinized tube for plasma insulin determination. Plasma insulin was measured using the Mouse Ultrasensitive Insulin ELISA kit according to the manufacturer’s instructions (ALPCO, Salem, NH, USA). HOMA-IR and HOMA-β were calculated based on glucose and insulin at 0 min.

### Tissue Collection

Mice were sacrificed at 16 weeks old. The body weight and organ weight were recorded. All collected samples were snap-frozen and stored at -80°C until further analysis. Cecal samples were collected and sent for shotgun metagenomic sequencing for a subset of mice (n=6 per group). Cecal samples were subject to shotgun sequencing and metabolomics analyses.

### Liver arsenic quantification

The analysis was performed by the Mass Spectrometry Core in the Research Resources Center of University of Illinois Chicago. Briefly, the mouse liver samples were prepared for analysis using the Perkin Elmer Sample Preparation Block. The samples were added to the digestion tubes. DI water, HNO_3_, and H_2_O_2_ were added to each digestion tube. The digestion tubes were placed in the Perkin Elmer Sample Preparation Block at 100°C for 130 minutes. After cooling, the samples were diluted with DI water. The IS working solution was added to the samples. All samples were automatically diluted with DI H_2_O to a final volume of 2× for analysis. All analyses were carried out on a Perkin Elmer NexION 2000 ICP-MS using the Elemental Scientific (ESI) prepFast automated dilution sample introduction system equipped with a SampleSense valve and peristaltic pump tubing. The samples were introduced at a flow rate of 300 μL/min using a PFA ST3 Type C nebulizer and glass cyclonic spray chamber maintained at 2°C. The RF power was set to 1600 W. Argon gas was used as plasma, auxiliary, and nebulizer gas with respective flow rates of 15.0 L/min, 1.2 L/min, and 0.98 L/min. MS data were acquired in KED mode with Helium as the cell gas at a flow rate of 4.8 mL/min.

### Metabolomics

All metabolomics analyses were performed at the Host-Microbiome Metabolomics Facility at the University of Chicago. Extraction solvent (80% methanol spiked with internal standards and stored at -80 °C) was added at a ratio of 100 mg of material/mL of extraction solvent in beadruptor tubes (Fisherbrand; 15-340-154). Samples were homogenized at 4 °C on a Bead Mill 24 Homogenizer (Fisher; 15-340-163), set at 1.6 m/s with 6 thirty-second cycles, 5 seconds off per cycle. Samples were then centrifuged at -10 °C at 20,000 x g for 15 min to generate supernatants for subsequent metabolomic analysis. All samples are extracted using a solvent containing isotopically labeled internal standards with known concentrations to assess metabolite extraction efficiency and instrument performance.

Gas chromatography-mass spectrometry (GC-MS) was used to detect 49 compounds following derivatization with pentafluorobenzyl bromide (PFBBr)(Haak et al. 2018). SCFAs (acetate, butyrate, propionate) as well as 5-aminovalerate, glycine, proline, succinate, and tyramine were quantitatively analyzed following PFB derivatization and detection by negative collision-induced gas chromatography-mass spectrometry ((-)CI-GC-MS, Agilent 8890). These, along with an additional 41 compounds, are reported as normalized relative abundances and include tryptophan catabolites, indoles, amino acids, branched-chain fatty acids, phenolic and aromatic compounds. Sixty-nine bile acids from the primary, secondary, and glyco/tauro-conjugated subclasses were analyzed using negative mode liquid chromatography-electrospray ionization-quadrupole time-of-flight-MS ((-)LC-ESI-QTOF-MS, Agilent 6546)(Gómez et al. 2020). In addition to retention time validation, the standard intact and fragment masses are routinely detected with <5 ppm error relative to calculated values.

### Statistical analyses

Data were analyzed in a 2 × 2 design crossing facility (InstA, InstB) with drinking-water treatment (Control or iAs), producing four groups: InstA.Control, InstA.iAs, InstB.Control, and InstB.iAs. All statistical analyses were conducted in R (v. 4.5.0). Univariate testing was performed with the rstatix package (v. 0.7.3), and figures were generated with tidyverse packages (v.2.0.0).

Glucose-tolerance-test area under the curve (GTT AUC), homeostatic model assessment of insulin resistance and β-cell function (HOMA-IR and HOMA-β, respectively), fasting glucose, fasting insulin, pancreas weight (% body weight), body weight, liver arsenic, and liver trace metals were tested for normality within each group using a Shapiro-Wilk test. Pairwise comparisons among the four groups used Welch *t*-tests or Wilcoxon rank-sum tests. Pairwise comparisons were calculated within institutions (Control v. iAs), between institutions’ control groups, or between institutions’ iAs groups; no calculation was made between institutions’ iAs and control groups (*e.g.,* InstA.Control v. InstB.iAs). A two-way aligned rank transform (ART) ANOVA (ARTool v0.11.2) was used as a nonparametric factorial test of the facility × iAs-treatment interaction for GTT-AUC, HOMA-IR, HOMA-β, fasting glucose, fasting insulin, pancreas mass (% body weight), body weight, and body-weight change.

For multivariate ordination of targeted and untargeted metabolites, the normalized metabolite data were used. Any metabolite with one or more missing values was removed. Features were mean-centered and scaled to unit variance within the principal components analysis (prcomp). To identify the metabolites contributing most to the ordination, the five features with the largest combined PC1/PC2 loading magnitude were overlaid as biplot vectors. Group separation along the ordination was tested by permutational multivariate analysis of variance (PERMANOVA) on Euclidean distances of the scaled feature matrix using the Vegan package (v.2.7-5) with location and treatment as factors and 10,000 permutations. For univariate analysis of targeted metabolites, analyses were performed on the quantified subset of metabolites against reference standards. Each metabolite was screened for a treatment or facility (location) effect using Wilcoxon rank-sum tests.

### Taxonomic analyses

Metagenomic sequencing reads were processed with a custom pipeline described previously (Metwally et al. 2020) hosted on the University of Illinois Chicago high-performance computer cluster “Extreme.” Briefly, low-quality reads (<25 Phred quality score), short reads (<100 bp), and human reads were filtered. Short reads were then assembled using MetaVelvet (Namiki et al. 2012). Taxonomic profiles for each sample were constructed using WEVOTE (Metwally et al. 2016), with Kraken (Wood, Lu, and Langmead 2019), Clark (Ounit et al. 2015), and BLASTN (Altschul et al. 1990), as base classifiers.

Following taxonomic assignment, taxonomic profiles were imported into R (v 4.5.0) and processed using the package phyloseq (v 54.1; McMurdie & Holmes, 2013). Compositional structure (beta-diversity) was assessed by PCA of centered-log-ratio (CLR)-transformed species abundances. A PERMANOVA test was conducted within the R package Vegan (v.2.7-5). PERMANOVA tests and p-values were calculated based on 10,000 permutations. Alpha diversity measures were generated using the ‘microbiome’ package (v1.32.0; Lahti & Shetty, 2017), computed on rarefied data due to differences in sequencing depth between institutions. Richness and diversity indices (observed richness, Shannon, inverse Simpson, Gini-Simpson, Fisher) and evenness metrics (Pielou, Bulla, Camargo, eVar, Simpson, coverage) between groups were compared by Wilcoxon tests.

Per-taxon negative-binomial generalized linear models using the MASS package (v 7.3-65) were fit with terms for facility (InstA vs InstB), treatment (iAs vs control), and their interaction. A model was fit for every taxon present in at least 75% of samples. To account for differences in sequencing depth across samples, each model included an offset term of log-transformed sample read count. Statistical significance of log2 fold-change coefficients was assessed, and p-values were adjusted for false discovery rate using the Benjamini–Hochberg procedure.

Several random forest (RF) models were used with taxon abundances as features for the prediction of outcome variables. Taxon abundances were filtered by mean relative abundance, and a random forest was generated for each abundance threshold: 1% (22 taxa), 0.1% (96 taxa), and 0.01% (749 taxa) from an original set of 3707 taxa to reduce the number of predictors. For each model, variable importance was taken from a fit on all samples, and prediction error was estimated separately using leave-one-out cross-validation (LOOCV), where the model was retrained on n−1 samples and used to predict the held-out sample across all folds. RF classifiers (1,001 trees) were trained to predict facility, treatment, and facility x treatment interaction. Out-of-bag (OOB) error from the full-data fit, pooled LOOCV accuracy relative to the majority-class baseline, and variable importance (mean decrease in accuracy) were reported.

To assess associations between gut microbial taxa and quantified targeted metabolites while eliminating the confounding influence of housing facility, correlations were computed separately within each facility (n = 12 mice). Taxa with a mean relative abundance ≥ 1% within that facility were retained and CLR-transformed. Each retained taxon was correlated against each metabolite across the facility’s mice using Spearman’s rank correlation. Pairs with fewer than six observations, or with zero variance in either variable, were not tested. Within each facility, p-values were corrected for multiple comparisons across taxa within each metabolite using the Benjamini–Hochberg procedure. Nominal significance was defined as p < 0.05 and false-discovery-rate significance as q < 0.05.

### Metagenome functional analyses

Raw paired-end reads from 24 samples were quality-trimmed and filtered using the read QC module of MetaWRAP (v1.3.2) with the –skip-bmtagger flag (Uritskiy, DiRuggiero, and Taylor 2018). Host (mouse) sequences were removed by mapping the quality-filtered reads against the *Mus musculus* reference genome (GRCm39, Ensembl release 115) with Bowtie2 under –very-sensitive-local settings (Langmead and Salzberg 2012). Host-filtered reads were assembled per sample using metaSPAdes via the MetaWRAP assembly module, retaining contigs ≥500 bp (Bankevich et al. 2012). To improve recovery of facility-associated taxa and given the similarity in taxonomic composition within facilities as determined by taxonomic analysis, per-facility co-assemblies were also generated by pooling host-filtered reads from all twelve samples within each facility and assembling each pool with MEGAHIT (Li et al. 2015), retaining contigs ≥300 bp.

#### Per facility gene catalog

Protein-coding genes were predicted from all individual-sample assemblies and per-facility co-assemblies using Prodigal in metagenomic mode (-p meta)(Hyatt et al. 2010). A non-redundant gene catalog was constructed by two-stage clustering with CD-HIT (Fu et al. 2012). Predicted proteins from the individual assemblies were first clustered at 95% sequence identity with 90% alignment coverage of the shorter sequence (-c 0.95 -aS 0.9). The resulting representatives were then combined with the co-assembly proteins and re-clustered under the same thresholds. The final catalog comprised 1,487,379 non-redundant protein sequences, with matching nucleotide sequences extracted for read mapping. A Bowtie2 index was built from the non-redundant nucleotide catalog, and host-filtered reads from each sample were mapped back to the catalog under –very-sensitive –no-unal. Alignments were sorted and indexed with SAMtools (Danecek et al. 2021). Per-gene abundances were quantified with CoverM (contig mode) as read counts, TPM, and gene length, retaining only properly paired alignments with ≥95% read identity and ≥75% read alignment (--contig-end-exclusion 0) (Aroney et al. 2025). Differentially abundant genes per group were calculated using DESeq2 (Love, Huber, and Anders 2014), and only genes with log-fold-changes of at least >1 or <-1 were considered differentially abundant.

Arsenic-cycling genes were identified using AsgeneDB, a manually curated arsenic metabolism gene database comprising 400k + representative sequences from 59 arsenic metabolism gene families (Song et al. 2022). Gene catalog protein sequences were mapped against AsgeneDB using DIAMOND blastp (e-value ≤1e-10, ≥80% query coverage, ≥50% subject coverage) (Buchfink, Xie, and Huson 2015; Song et al. 2022). The top 10 hits were sorted by bit-score, e-value, and percent identity to select the top hit, and only annotations with >80% identity were retained. Carbohydrate-active enzymes were annotated with run_dbcan in protein mode, and genes annotated with at least 2 tools were retained (Zheng et al. 2023). General orthology and pathway annotation (COG, KEGG) was performed with eggNOG-mapper in DIAMOND mode against the eggNOG 5.0 database (emapperdb-5.0.2)(Huerta-Cepas et al. 2019).

#### Metagenome-assembled genomes (MAGs)

MAGs were recovered from each per-facility co-assembly. Initial binning was performed with the MetaWRAP binning module using MetaBAT2, MaxBin2, and CONCOCT(Alneberg et al. 2014; Kang et al. 2015; Wu, Simmons, and Singer 2016), with sample reads supplied per facility. Bin sets were consolidated with the MetaWRAP bin_refinement module, retaining MAGs with ≥90% completeness and ≤5% contamination as estimated by CheckM (-c 90 -x 5)(Parks et al. 2015). Refined MAGs were quantified across samples in MetaWRAP with quant_bins, taxonomically classified with classify_bins, and functionally annotated with annotate_bins. Arsenic-cycling genes and CAZymes were additionally annotated per MAG using DIAMOND against AsgeneDB and run_dbcan, respectively, as described for the gene catalogs.

## Results

### Housing facility dominates metabolic phenotype, with facility-specific arsenic effects

Mice were housed at one of two facilities and received either 50 ppm inorganic arsenic (iAs) in drinking water or control water *ad libitum* to produce four groups: InstA.Control, InstB.iAs, InstA.Control, InstB.iAs. Within the control groups, InstB animals were consistently more glucose-intolerant than InstA animals. GTT AUC was higher in InstB than InstA across both control and iAs treatment groups (Fig. 1B). HOMA-IR, fasting glucose, and fasting insulin were elevated in InstB.Control compared to InstA.Control at 8 weeks and remained higher at 12 weeks (Fig 1B; E-F). InstB.Control mice were also consistently higher in weight from the start of the exposure experiment and gained more weight over time (Fig. 2B-C). Relative pancreas weight (as % body weight) was roughly two-fold greater in InstB than InstA (p = 2.1e-5) across the control groups.

**Figure 1.**
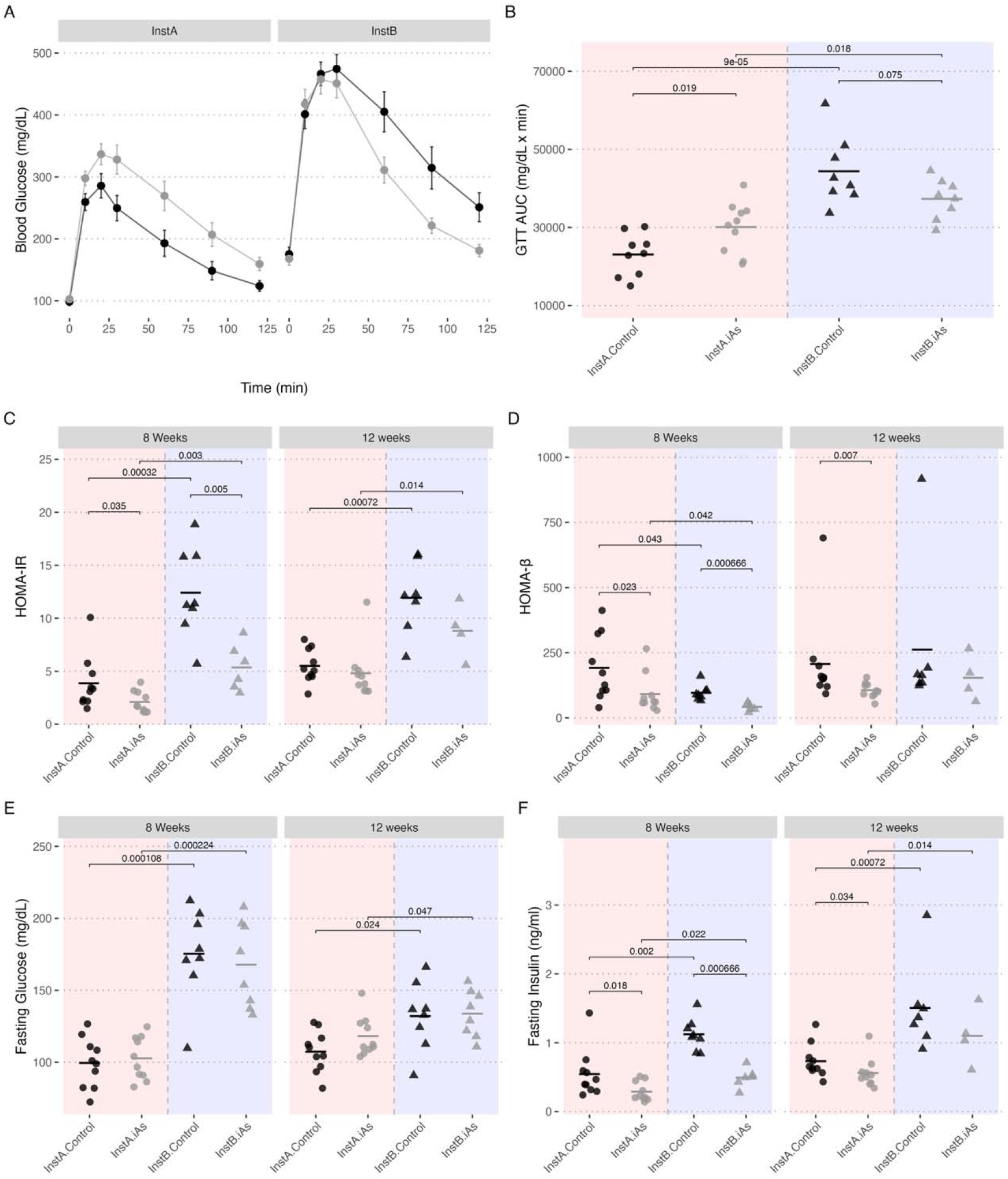
**(A)** Glucose tolerance test (GTT) time course at 8 weeks of exposure, shown as mean blood glucose ± SEM at each time point. **(B)** Area under the GTT curve (AUC) at 8 weeks. **(C)** HOMA-IR, **(D)** HOMA-β, **(E)** fasting glucose, and **(F)** fasting insulin, each at 8 and 12 weeks of exposure. Group comparisons used Welch’s t-test (B, E, F) or the Wilcoxon rank-sum test (C, D), selected by Shapiro–Wilk normality testing; only comparisons where p < 0.1 are displayed. Horizontal crossbars indicate group means.

**Figure 2.**
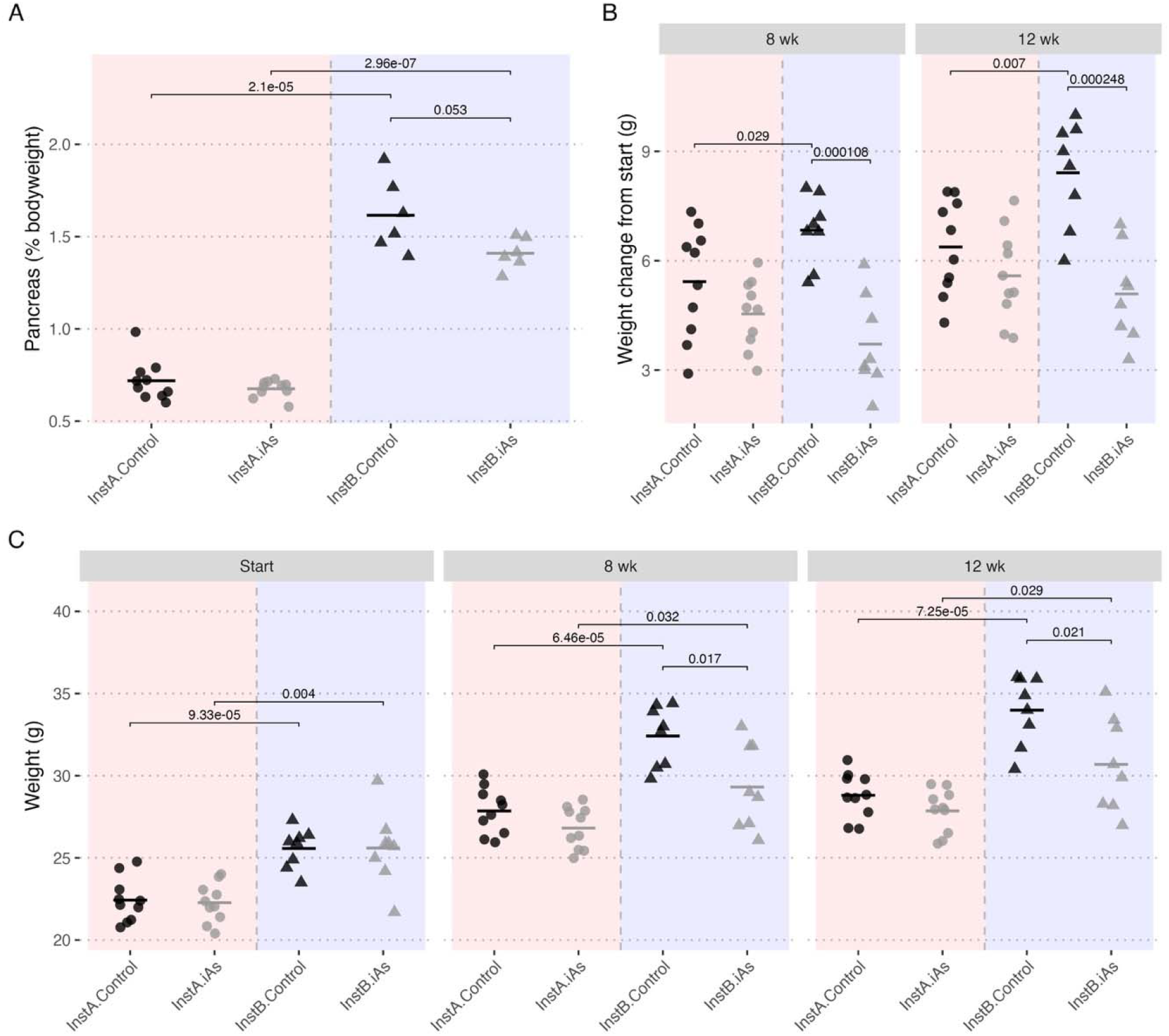
**(A)** Pancreas weight as a percentage of bodyweight. **(B)** Body weight change from the start of exposure treatment. **(C)** Body weight at study start and after 8 and 12 weeks of exposure. Groups were compared by Welch’s t-test; only comparisons where p < 0.1 are displayed.

Despite baseline differences, iAs exposure produced facility-dependent effects on metabolic phenotypes. Notably, in InstA, arsenic impaired the ability to clear glucose (InstA.Control vs InstA.iAs p = 0.019). Whereas in InstB, the effect trended in the opposite direction (InstB.Control vs InstB.iAs p = 0.075), with lower average GTT AUC in the iAs group at InstB. Within InsB, iAs-treated mice weighed less than controls at both 8 and 12 weeks (p = 0.017 and 0.021), an effect absent in InstA (p = 0.102 and 0.135). ART ANOVA tests indicated significant facility x iAs interaction terms for GTT AUC (p = 0.004), fasting insulin at 8 weeks (p = 0.002), HOMA-IR at 8 weeks (p =0.003), and relative pancreas weight (p =0.003). Body weight change interaction terms were significant both at 8 (p=0.009) and 12 weeks (p =0.034). No other phenotypic endpoints indicated significant facility x iAs interactions at the 12-week time point.

Hepatic arsenic confirmed successful exposure at both sites (Fig. 3). Arsenic treatment raised liver arsenic in both facilities, with no significant differences between iAs treatment groups (p = 0.637), although InstB hepatic arsenic was more variable among animals. Control mice were indistinguishable across facilities (p = 0.306), confirming near-zero baseline arsenic exposure at both sites. Other hepatic trace metals (SuppFig. 1) differed by facility (Co, Cu, Mn, and Mo, p ≤ 0.05) but showed no significant arsenic-mediated effects within facility, indicating source water within each facility carried significantly different trace metal mixtures despite originating from the same municipal water source. Animals with greater tissue arsenic accumulation trended towards more weight gained by 12 weeks, but the association did not reach significance (Fig. 4B, p = 0.079). Hepatic arsenic was not significantly associated with other metabolic outcomes in the iAs treatment groups (SuppFig2).

**Figure 3.**
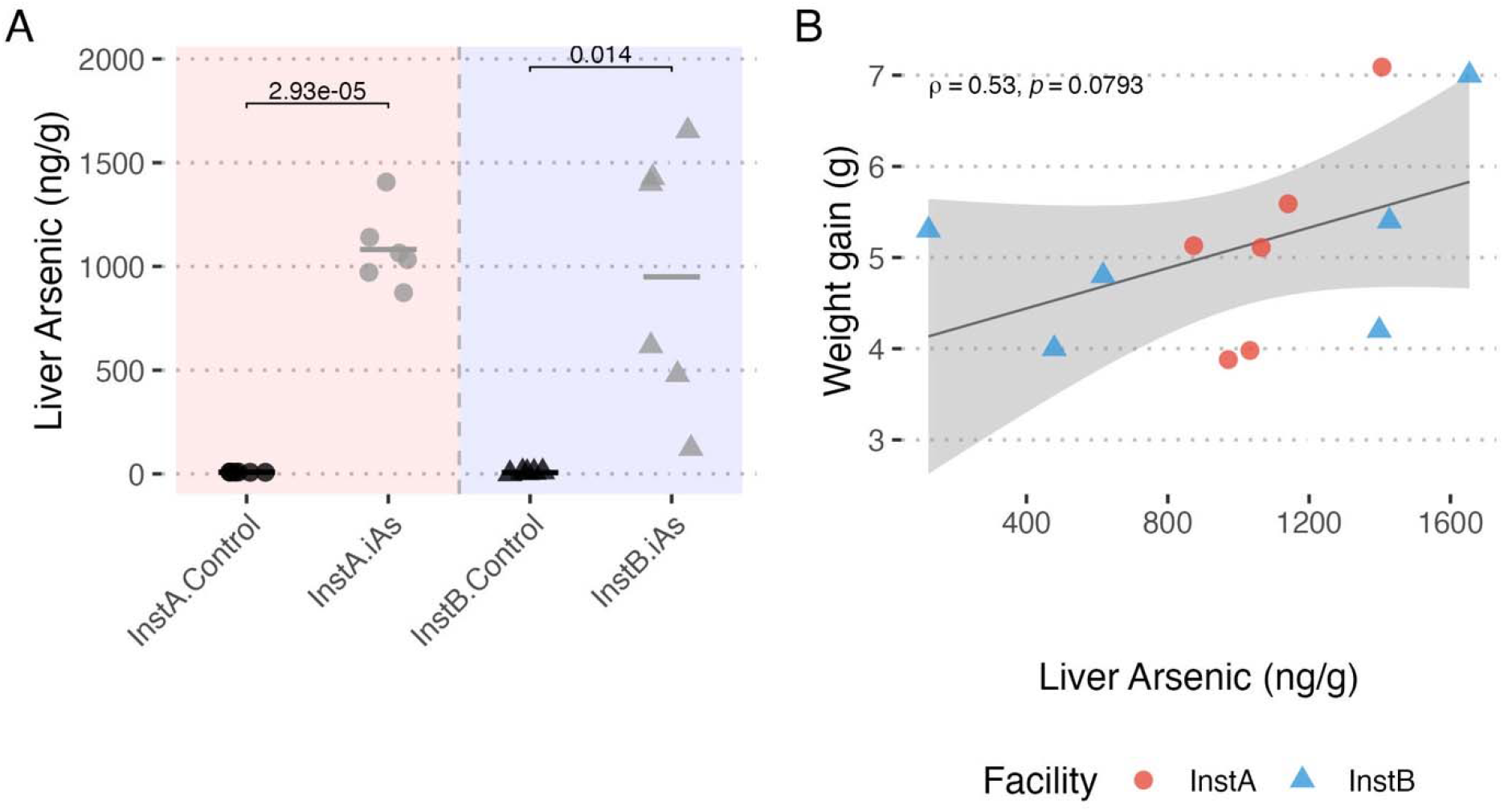
**A)** Liver arsenic concentration (ng/g) by group. Groups were compared by Welch’s t-test; only comparisons where p < 0.1 are displayed. **B)** Liver arsenic in the iAs-treated mice compared to weight gain at 12 weeks.

**Figure 4.**
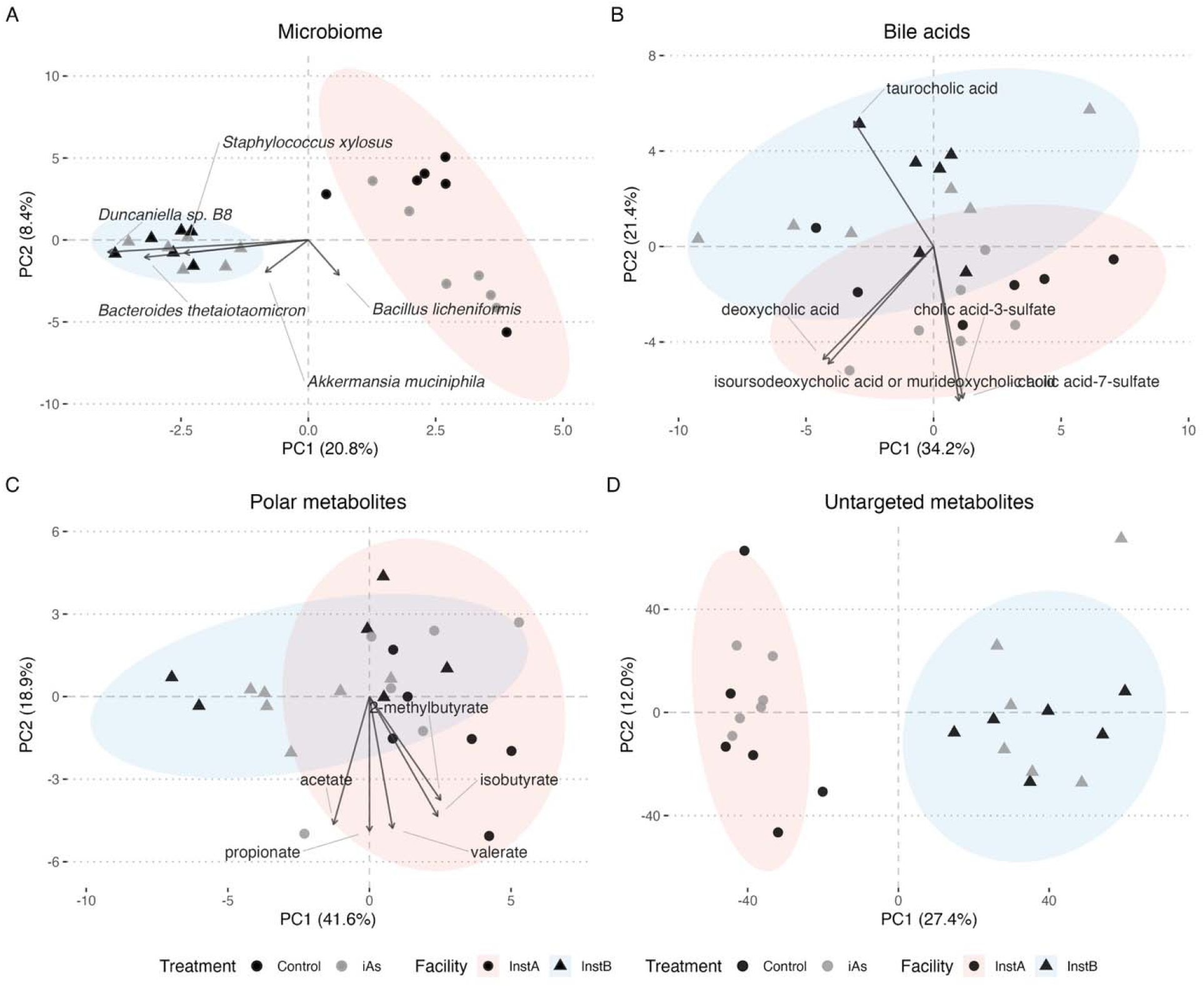
Principal components analysis of (A) CLR-transformed microbial species abundances (Aitchison distance), (B) targeted bile acids, (C) targeted polar metabolites, and (D) untargeted metabolites. Shaded ellipses are 95% confidence regions by facility (InstA, red; InstB, blue). In panels A-C, gray arrows show the five features with the largest combined PC1/PC2 loadings, scaled to the score range and labeled with the corresponding taxon or metabolite.

### Facility structures the multi-omic landscape

PCA on microbial species, targeted metabolites, and untargeted metabolites revealed strong facility-level separation that overprinted iAs treatment effects (Fig. 4). PCA of CLR-transformed microbiome profiles separated InstA from InstB along PC1 (20.8% of variance), with *Bacillus licheniformis* loading toward InstA, and *Bacteroides thetaiotaomicron*, *Duncaniella sp. B8*, *Staphylococcus xylosus*, and *Akkermansia muciniphila* loading toward InstB. The same facility split was evident for untargeted metabolites (PC1 27.4%, PC2 12.0%), with facility resolving along PC1. Bile acids (PC1 34.2%, PC2 21.4%) and polar metabolites (PC1 41.6%, PC2 18.9%) displayed slightly more overlap for facility along PC1 and PC2. In the bile acid ordination, sulfated species (cholic acid-3-sulfate, cholic acid-7-sulfate) loaded toward InstA, while taurocholic acid loaded toward InstB. In the polar metabolite ordination, the short-chain fatty acids (acetate, propionate, isobutyrate, 2-methylbutyrate, valerate) loaded together toward InstA. The top 5 loadings for the untargeted metabolites were unannotated.

PERMANOVA on the same features confirmed facility as the dominant axis of variation across every data layer (Table 1). Location explained a substantial and significant fraction of variance in the microbiome (R² = 0.19, p = 0.0001), bile acids (R² = 0.19, p = 0.0002), polar metabolites (R² = 0.20, p = 0.0005), and untargeted metabolites (R² = 0.26, p = 0.0001). In contrast, iAs treatment explained little variance and did not reach significance in any of the four layers.

**Table 1:** PERMANOVA of microbiome (Aitchison distance of taxon abundances) and metabolite ordinations (Euclidean distance on scaled features).

| Term | Df | R<sup>2</sup> | pseudo-F | p |
| --- | --- | --- | --- | --- |
| **Microbiome** |  |  |  |  |
| Facility | 1 | 0.193 | 5.270 | **0.0001** |
| Treatment | 1 | 0.041 | 0.939 | 0.4736 |
| **Bile acids** |  |  |  |  |
| Facility | 1 | 0.187 | 5.061 | **0.0001** |
| Treatment | 1 | 0.036 | 0.825 | 0.5530 |
| **Polar metabolites** |  |  |  |  |
| Facility | 1 | 0.201 | 5.538 | **0.0001** |
| Treatment | 1 | 0.027 | 0.610 | 0.7266 |
| **Untargeted metabolites** |  |  |  |  |
| Facility | 1 | 0.257 | 7.592 | **0.0001** |
| Treatment | 1 | 0.049 | 1.142 | 0.2458 |

Pairwise Wilcoxon-rank sum testing on quantified metabolites (polar metabolites and bile acids) indicated that none of the quantified metabolites were significantly different within institutions between treatment groups. Taurocholic acid, a top loading in the bile acid ordination, was elevated at InstB relative to InstA, with no separation between Control and iAs within either facility (SuppFig3). Among the polar metabolites, butyrate was higher at InstB in the iAs treatment group compared to InstA, whereas glycine showed the opposite trend (SuppFig4).

### Microbiome composition is facility-specific

A negative binomial generalized linear model (NB-GLM) was fit with terms for facility (InstA vs InstB), treatment (iAs vs control), and their interaction for each of 941 taxa at the species level that were present in at least 75% of samples. Taxa with absolute log2-fold change values of at least 1 were considered differentially abundant. Of the taxa modeled, 120 location terms, 30 iAs treatment terms, and 26 interaction terms were significant after Benjamini-Hochberg false discovery rate adjustment (SuppFig5). *B. thetaiotaomicron* showed the strongest site association, with 212.7-fold higher abundance at InstB than InstA (95% CI 152.6–296-fold; Fig 5). Six *Alistipes* species were also enriched at InstB relative to InstA: *A. dispar* (21.1-fold, CI 11.3–40), *A. shahii* (17.7-fold, CI 10.3–30), *A. finegoldii* (14.0-fold, CI 8.2–24), *A. onderdonkii* (13.1-fold, CI 7.5–23), *A. communis* (12.9-fold, CI 7.7–22), and *A. megaguti* (11.2-fold, CI 6.7–19). Several species were enriched at InstA, including *Sarcina* sp. JB2 (27.1-fold lower at InstB, CI 9.4–78.2-fold), *Lactobacillus iners* (11.8-fold lower, CI 4.2–33.1-fold), and *Clostridium chauvoei* (8.2-fold lower, CI 6.3–10.6-fold).

**Figure 5.**
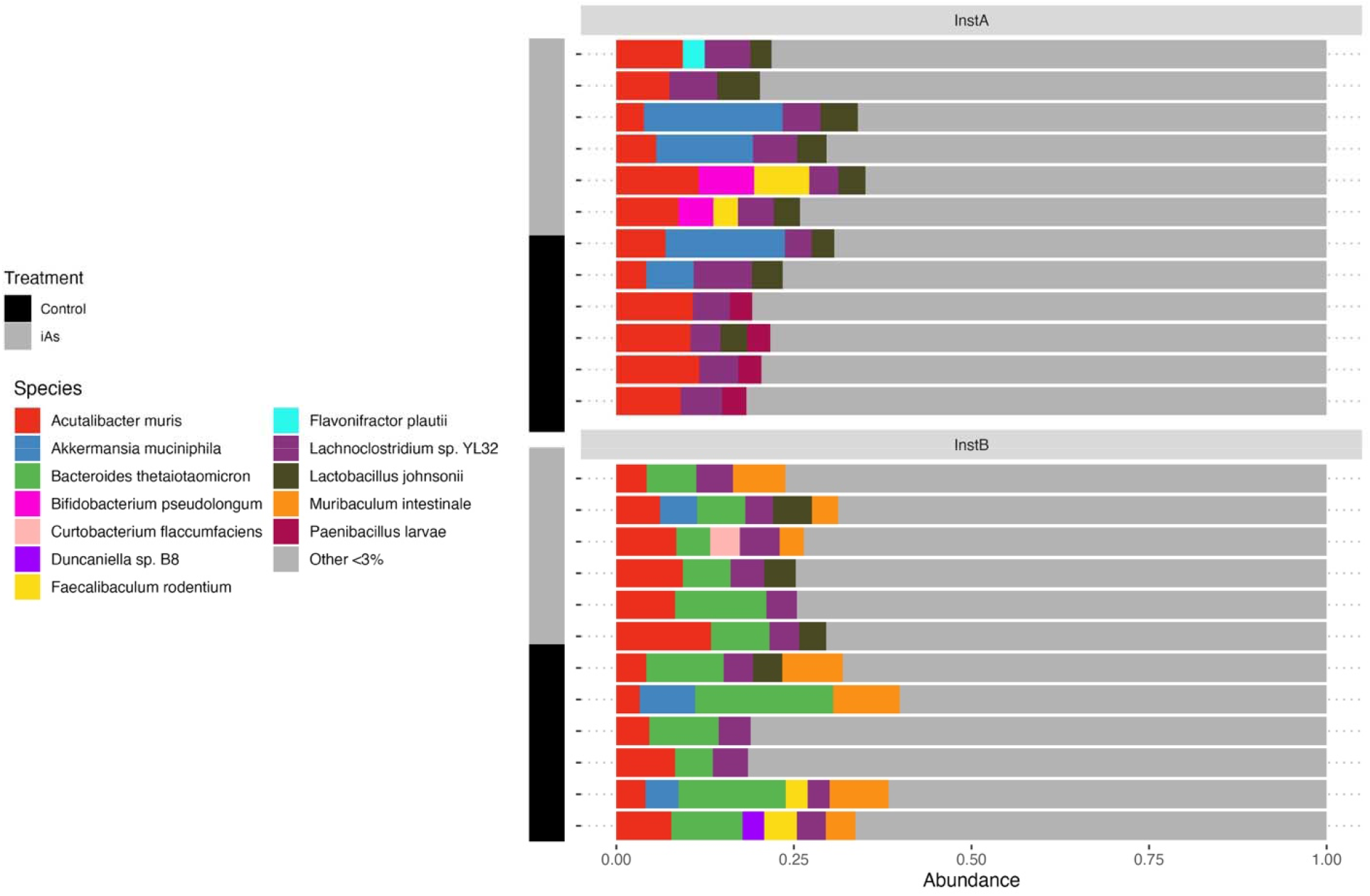
Gut microbiome species composition. Stacked bars of relative abundance of the most abundant species per animal, faceted by facility and annotated by treatment (Control, iAs); taxa below 3% are pooled as “Other.”

Arsenic exposure was associated with large shifts in several taxa at InstA, including *Bifidobacterium pseudolongum,* which increased 188.8-fold with iAs exposure (CI 26.6–1339-fold), and *Faecalibaculum rodentium,* which increased 16.0-fold (CI 3.2–80-fold). *Turicibacter sanguinis* (13.0-fold lower, CI 4.2–40.8-fold) and *Amycolatopsis mediterranei* (10.7-fold lower, CI 2.1–54.9-fold) decreased with iAs exposure. Several taxa showed a significant interaction effect, including some that were iAs-responsive at InstA, indicating the iAs response itself depended on facility. Most notably, when accounting for the interaction term, both *B. pseudolongum* and *F. rodentium* decreased in response to iAs at InstB, indicating an opposite iAs effect at InstB compared with InstA.

RF classification models were trained to predict facility, treatment, and group (facility + treatment) using taxon abundances that were filtered by mean relative abundance at multiple thresholds: 1% (22 taxa), 0.1% (96 taxa), and 0.01% (749 taxa; Table 2). Facility classification was highly accurate (96% LOOCV accuracy) with just the 22 most abundant taxa. LOOCV accuracy improved to 100% when 749 taxa were included in the model. Neither treatment nor interaction outcomes could be predicted to the same accuracy as facility alone. LOOCV accuracy for treatment and interaction models at all filter thresholds ranged between 54 and 67%. At the 1% mean abundance filter level, *B. thetaiotaomicron, Duncaniella sp.B8, Clostridioides difficile, Enterocloster bolteae,* and *A. muciniphila* were the most important predictors of facility (Supp. Fig 6).

**Table 2.** Random forest performance across relative abundance filters.

| Model | Filter | n taxa | OOB error | LOOCV accuracy | Baseline |
| --- | --- | --- | --- | --- | --- |
| Group (facility × treatment) | 1% | 22 | 0.458 | 0.542 | 0.25 |
| Group (facility × treatment) | 0.1% | 96 | 0.458 | 0.542 | 0.25 |
| Group (facility × treatment) | 0.01% | 749 | 0.458 | 0.625 | 0.25 |
| Facility | 1% | 22 | 0.042 | 0.958 | 0.50 |
| Facility | 0.1% | 96 | 0.042 | 0.958 | 0.50 |
| Facility | 0.01% | 749 | 0.000 | 1.000 | 0.50 |
| Treatment | 1% | 22 | 0.333 | 0.667 | 0.50 |
| Treatment | 0.1% | 96 | 0.417 | 0.542 | 0.50 |
| Treatment | 0.01% | 749 | 0.333 | 0.625 | 0.50 |

Unique taxa from the NB-GLMs that were significantly differentially abundant (159 total taxa) were further investigated to determine whether microbiome composition was associated with mouse phenotypes. Phenotypes tested were GTT-AUC at 8 weeks, log-transformed HOMA-IR*, HOMA-β*, fasting glucose, fasting insulin, and body-weight gain, all at 8 weeks and 12 weeks. The reduced community of 159 taxa was CLR-transformed and aggregated by taking the mean CLR value across ASVs sharing the same species label. A constrained ordination analysis on each phenotype using the CLR-transformed Aitchison distances was used to calculate the amount of phenotypic variation that could be explained by taxonomic composition. Significance was then assessed with a PERMANOVA test with 10000 permutations. GTT-AUC (R^2^ = 0.17, p = 0.0017), fasting insulin (R^2^ = 0.12, p = 0.013), log(HOMA-IR) (R^2^ = 0.18, p = 0.0017), and fasting glucose (R^2^ = 0.27, p = 0.00009) were significantly associated with the microbiome composition at 8 weeks. Similarly, at 12 weeks, fasting insulin (R^2^ = 0.20, p = 0.0003), log(HOMA-IR) (R^2^ = 0.23, p = 0.0009), and fasting glucose (R^2^ = 0.12, p = 0.01) were significantly associated with the microbiome composition at 12 weeks. *HOMA-β* and body-weight gain failed to reach significance during either time point.

Phenotypes with significant correlations to microbiome composition were further investigated by calculating the Spearman’s rank correlation for each individual taxon of the 159 total to each individual phenotype at 8 and 12 weeks. Phenotype x taxon associations were tested separately per facility using all 12 mice, regardless of treatment group. Because phenotypic associations are confounded by facility abundances of the taxa, we focused on associations that increased or decreased in the same direction at both facilities in response to phenotype within the control groups. Overall, 1,113 taxon x phenotype associations were tested, and 533 (48%) showed correlation of the same sign at each facility. When considering correlations that were significantly associated with phenotype, only one same-direction association was significant at both facilities; *Campylobacter coli* was positively associated with fasting glucose at InstA (ρ = 0.7) and InstB (ρ = 0.7). These results indicate that taxon x phenotype associations were highly facility-specific and phenotypic variation between facilities is likely the result of the overall community rather than driven by a single taxon.

Microbial alpha diversity richness and diversity indices (observed richness, Shannon, inverse Simpson, Gini-Simpson, Fisher) and evenness metrics (Pielou, Bulla, Camargo, eVar, Simpson, coverage) between groups were compared by Wilcoxon tests. Overall, alpha diversity was heterogeneous between samples of the same group, and most indices did not distinguish between the facility or treatment group. Significant differences were confined to Shannon diversity and to Bulla and Evar evenness, each of which was lower in the InstB control group relative to the InstA control (Suppl. Fig. 7).

### Metagenomic gene and functional content tracks facility

The gene catalog produced a total of 1,487,379 non-redundant genes across facilities. Differential abundance testing was conducted on genes with at least 10 counts in at least 4 samples, producing 18,628 genes tested. There were 11,733 (63%) differentially abundant genes between facilities at BH-adjusted p < 0.05 and log2 fold-change| > 1. Of those, 8,708 genes were enriched in InstB compared to 3,025 genes enriched in InstA (Fig. 6A). In contrast, only 139 genes were significantly differentially abundant in the iAs treatment groups. Arsenic metabolism genes were identified using AsgeneDB, a manually curated arsenic metabolism gene database (Song et al. 2022). The AsgeneDB contains 59 As metabolic gene families and includes metabolic pathways for transport, respiration, reduction, oxidation, and methylation/demethylation processes. Several arsenic metabolism genes were enriched in InstB relative to InstA, including *PiT* (transport)*, arsB* (transport)*, arsR* (reduction)*, pstC* (transport)*, and pstB* (transport). The arsenic transport genes labeled *PiT* were enriched by over 23 log2-fold change at InstB (Fig 6B). Hierarchical clustering of normalized arsenic metabolism gene counts revealed arsenic metabolism gene clusters by facility, but not arsenic treatment (Fig. 6C). Carbohydrate active (CAZyme) gene counts also clustered by facility (Supp Fig. 8), but not arsenic treatment.

**Figure 6.**
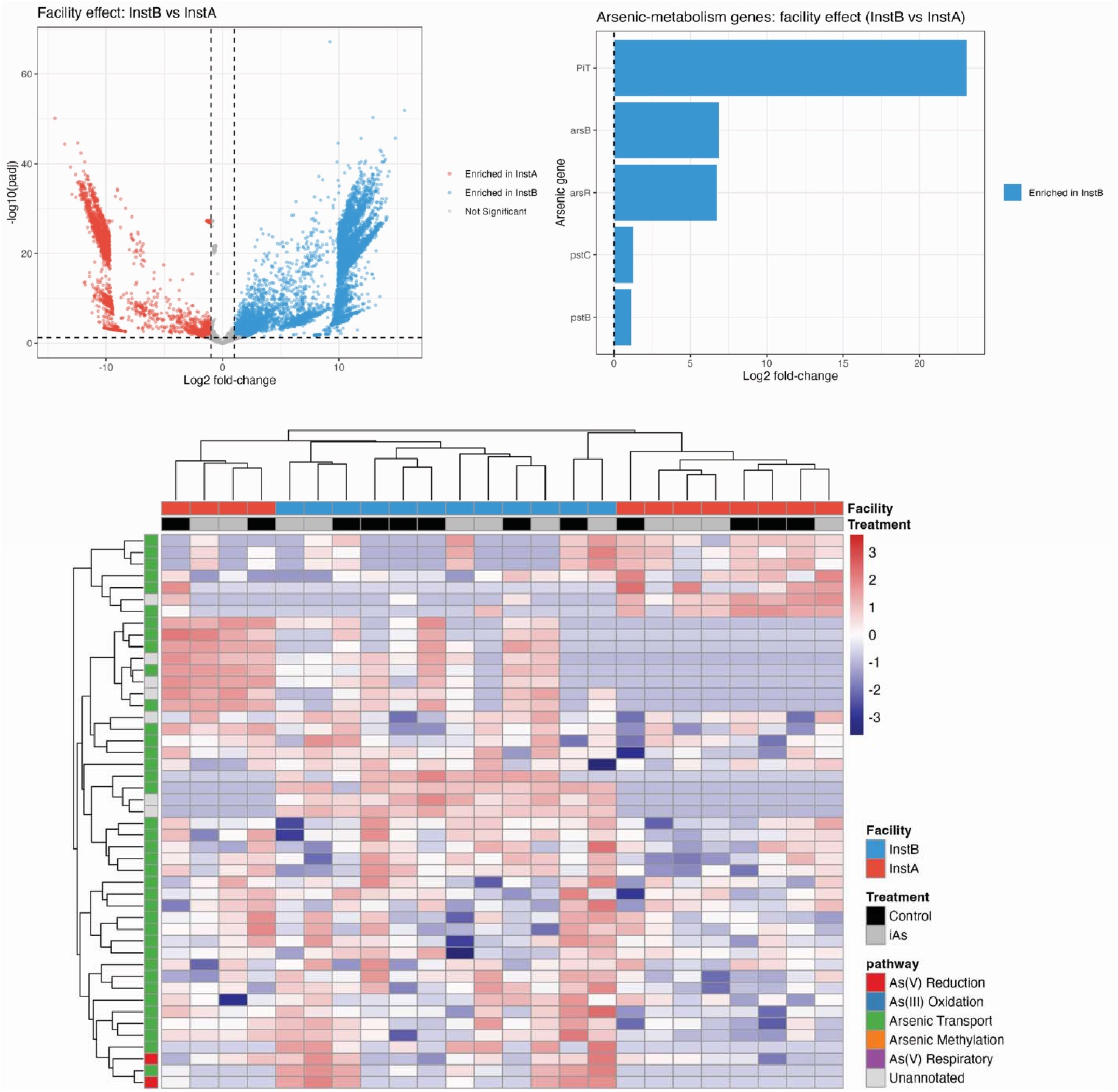
**A)** Volcano plot of gene-level differential abundance calculated using DESeq2, InstB vs InstA (red enriched in InstA, blue enriched in InstB; dashed lines mark adjusted p = 0.05 and |log2 fold-change| = 1). **B)** Differentially abundant arsenic metabolism genes across facilities. **C)** Hierarchically clustered heatmap of arsenic-metabolism gene abundances across samples. Cell color encodes the row-wise z-score of DESeq2-normalized counts.

None of the 139 genes that were differentially abundant in the iAs treatment groups were annotated as arsenic metabolism genes. Enrichment and depletion were roughly balanced in number, but the largest-magnitude shifts were depletions (Supp. Fig 9). The single most depleted features were DDE-family transposases (log₂FC −5), followed by a DNA-dependent RNA polymerase subunit (log₂FC −4) and methyl-accepting chemotaxis proteins (log₂FC −3.3). Genes enriched under iAs were dominated by carbohydrate-active and transport functions. Several glycan-processing families increased, including a glycosyl transferase group 1, glycoside hydrolase family 25, a depolymerase, and a 5′-nucleotidase.

MAGs were assembled per facility, and arsenic gene content per MAG was quantified (Supp. Fig. 10). Both facilities had similar arsenic cycling functional capability. At InstB, *B. thetaiotaomicron* possessed the greatest number of arsenic cycling genes, and at InstA, *B. pseudolongum* contained the greatest number of arsenic metabolism genes. Several MAGs were assembled separately at both facilities, including *Lactobacillus johnsonii, F. rodentium,* and *A. muciniphila*.

### Taxon–metabolite associations are facility-specific (**Fig. 7**)

Spearman rank correlations were computed for each housing facility separately between every taxon retained at ≥1% mean relative abundance within that facility and targeted metabolite. Analyzing each facility on its own removes between-site structure, so each coefficient reflects a within-cohort taxon–metabolite relationship. Both cohorts yielded a modest number of nominally significant correlations (19 pairs in InstA, 35 in InstB at p < 0.05).

**Figure 7.**
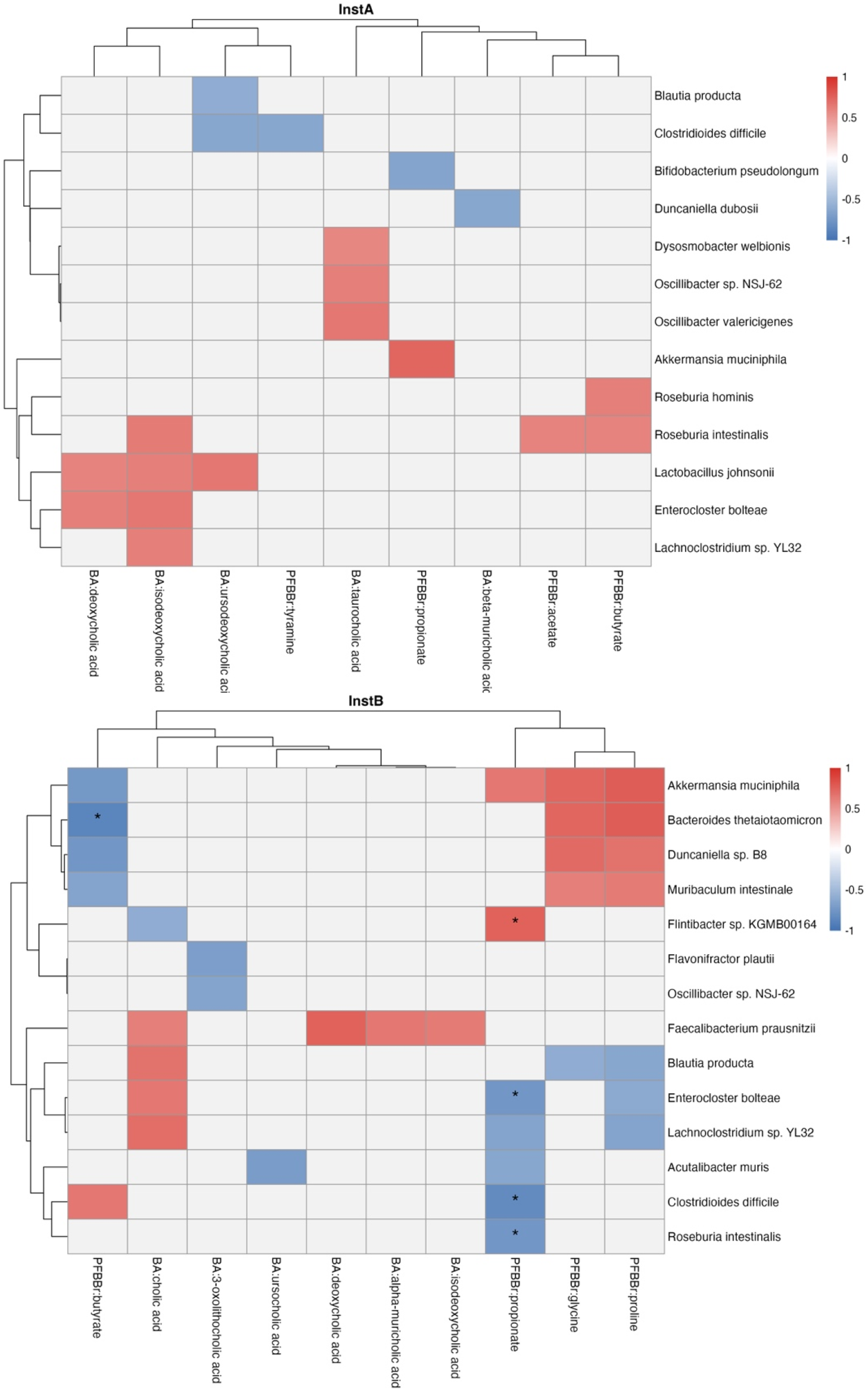
Within-facility associations between microbial taxa and targeted metabolites. Cell color shows the correlation coefficient (ρ) for pairs reaching nominal significance (p < 0.05); non-significant and untested pairs are left blank. Asterisks mark pairs surviving Benjamini–Hochberg correction (q < 0.05). Rows and columns are ordered by hierarchical clustering (Euclidean distance, complete linkage). Only taxa and metabolites with at least one nominally significant association are shown.

At InstA, the butyrate producers *Roseburia hominis* and *R. intestinalis* correlated positively with butyrate (and *R. intestinalis* additionally with acetate), and a cluster of *Lactobacillus johnsonii*, *Enterocloster bolteae*, and *Lachnoclostridium* sp. YL32 correlated positively with the unconjugated/secondary bile acids deoxycholic and isodeoxycholic acid. At InstB, *Faecalibacterium prausnitzii* correlated positively with several bile acids (cholic, deoxycholic, α-muricholic, and isodeoxycholic acid), and a co-clustering block of mucin- and Bacteroidetes-associated taxa (*Akkermansia muciniphila*, *B. thetaiotaomicron*, *Duncaniella* sp. B8, *Muribaculum intestinale*) shared a common signature of positive correlation with glycine and proline and negative correlation with butyrate. Signal surviving BH correction was sparse and confined entirely to InstB. Five taxon–metabolite pairs passed q < 0.05, and four of them involved a single metabolite, propionate: *Flintibacter* sp. KGMB00164 correlated positively with propionate, while *Enterocloster bolteae*, *C. difficile*, and *Roseburia intestinalis* correlated negatively.

The two facilities were largely non-overlapping in their association structure. For example, *R. intestinalis* correlated positively with SCFAs in InstA but negatively with propionate in InstB, and *C. difficile* associated with different metabolites in each cohort. The clearest directionally consistent exception was *A. muciniphila*, which correlated positively with propionate in both facilities.

## Discussion

The gut microbiome plays a key role in how environmental toxicants affect host health, serving as both a target and a mediator of exposure and toxicity. In this study, we examined how 50 ppm iAs affected the microbiome and metabolism of male C57BL/6J mice housed at two different facilities. We found that the facility, rather than iAs treatment, was the main source of variation across all measured data. Mice with the same genetics and iAs treatment showed differences in their microbiome, metabolites, glucose tolerance, body weight, and microbial functional gene content across facilities, while the effect of iAs was modest and facility-specific.

These results are consistent with previous work showing that the environment is a more important determinant of the gut microbiome than host genetics (Grieneisen et al. 2021; Rothschild et al. 2018). The two parallel facility experiments differed in source water, light cycle, and calendar time; therefore, the facility variable is confounded by several factors that cannot be resolved. However, this design captures the facility-level effects inherent in experimental rodent studies and addresses a well-recognized issue in microbiome research that treatment groups housed separately can produce apparent microbiome differences unrelated to the experimental intervention (Singh et al. 2021; Turner 2018; Voelkl et al. 2020).

In this work, the baseline microbial communities and metabolic phenotypes at InstA and InstB differ, and this difference propagates into the iAs treatment effect. Ordination analyses of species abundances and metabolomic profiles revealed facility-level separation along the first two principal components, while iAs treatment explained little variance and did not reach significance. PERMANOVA testing indicated facility-level differences explained between 19 and 25% of the variance between samples. In addition, a random-forest classifier predicted facility from taxon abundances with 96% LOOCV accuracy using only the 22 most abundant taxa, while the same accuracy could not be achieved for iAs treatment and group (treatment × facility), regardless of the number of species included in the model. Among untreated controls, InstB mice were consistently more glucose-intolerant than their InstA counterparts, with higher GTT area under the curve, elevated HOMA-IR, fasting glucose, and fasting insulin at both eight and twelve weeks, and higher body weights and weight gain over the course of treatment. Hepatic arsenic quantification nonetheless confirmed that iAs exposure was comparable across sites, with no significant differences in hepatic arsenic concentrations, although the InstB animals showed greater variability.

Despite the baseline differences between InstA and InstB animals, iAs treatment did produce a measurable facility-dependent effect. At InstA, arsenic significantly impaired the ability to clear a glucose load, whereas at InstB the effect trended in the opposite direction, with iAs-treated animals at InstB showing improved glucose tolerance. At InstB, iAs-treated animals also gained significantly less weight than controls. Arsenic is a well-documented suppressor of weight gain and adiposity, and arsenite can reduce adipocyte differentiation and impair adipose expansion (Carmean et al. 2020; Handali and Rezaei 2021; Trouba, Wauson, and Vorce 2000; Wauson, Langan, and Vorce 2002). An apparent improvement in glucose tolerance that coincides with reduced weight gain is therefore difficult to distinguish from the effects of weight loss. However, previous work has shown that exposure to iAs can worsen glucose handling while simultaneously being antiobesogenic (Carmean et al. 2020). In addition, in our data, hepatic arsenic concentrations showed a modest marginal positive correlation with weight gain (ρ r =0.53, p = 0.08), indicating glucose handling improvement by iAs-treated mice in InstB is likely the result of a mechanism independent of decreased adiposity.

Several taxa that responded to iAs treatment at InstA showed significant interaction terms, indicating that the direction of the arsenic response flipped at InstB, including *Bifidobacterium pseudolongum* and *Faecalibaculum rodentium*. *B. pseudolongum* has been reported as metabolically beneficial with the potential to treat diet-induced obesity (Bo et al. 2020), and has even been co-administered with arsenic trioxide to restore gut homeostasis and mitigate promyelocytic leukemia treatment-associated dysbiosis (Guo et al. 2026). However, in our dataset, it increased dramatically by ∼188-fold with iAs at InstA and was associated with worse glucose clearance, as it was positively associated with GTT AUC.

Our results suggest that host metabolic phenotype is determined by the overall configuration of the microbial community rather than by the abundance of a limited number of marker species. Constrained ordination demonstrated that a reduced set of 159 differentially abundant taxa identified by NB-GLM models accounted for a significant proportion of the variance in fasting insulin, HOMA-IR, GTT area under the curve, and fasting glucose. Despite these community-level associations, individual taxon–phenotype correlations were predominantly facility-specific. The single most site-discriminating taxon, *Bacteroides thetaiotaomicron,* was enriched over 200-fold at InstB. *B. thetaiotaomicron* has been shown to promote fat deposition and impair glucose tolerance in high fat diet fed mice (Cho, Cho, and Park 2022), potentially accounting for the baseline difference in metabolic phenotype at InstB.

The toxicity and mobility of arsenic depend on its chemical speciation, which is actively modified by both the host and the resident microbiota (McDermott et al., 2020). Functional metagenomics revealed gene-level differences that were mainly facility-specific. Differential abundance testing showed nearly two-thirds of the tested genes varied between facilities, compared to 139 differentially abundant treatment-responsive genes. Hierarchical clustering of both arsenic-metabolism genes and carbohydrate-active enzymes grouped samples by facility rather than by arsenic exposure. Both facilities nonetheless contained a broadly comparable repertoire of arsenic-cycling pathways and a similar arsenic-handling potential at the level of metagenome assembled genomes, indicating that the machinery for arsenic biotransformation was present in both communities even though the taxa carrying it differed. At InstB, the high-abundance *B. thetaiotaomicron* carried the most complete arsenic-gene pathways, compared to *B. pseudolongum* at InstA.

Short-chain fatty acids are strongly associated with host glucose homeostasis and insulin sensitivity, and reduced levels of SCFA-producing bacteria are associated with T2DM (Chong et al. 2024). Taxon–metabolite correlations were largely site-specific, with little shared association structure between the two cohorts, apart from a positive association of *A. muciniphila* with propionate at both sites, consistent with its established role as a mucin degrader and propionate producer that promotes glucagon-like peptide-1 secretion and improved glucose handling (Yoon et al., 2021). Butyrate was significantly higher in InstB.iAs than InstA.iAs (p = 0.002), yet this elevation could not be attributed to any single producer. No canonical butyrate-producing species was enriched at InstB. InstB was instead enriched for polysaccharide-degrading, acetate-producing Bacteroidota, including *B. thetaiotaomicron*, which are reported to support downstream butyrogenesis via cross-feeding (Chia et al. 2020). In our study, *B. thetaiotaomicron* was significantly negatively correlated with butyrate within InstB (q < 0.05), as were the other dominant Bacteroidota (*Duncaniella* sp. B8, *Muribaculum intestinale*). However, correlations were restricted to taxa exceeding 1% mean relative abundance, so a lower-abundance butyrogen expanding in the higher-butyrate animals would not have been tested, and its expansion would depress the CLR of the dominant degraders regardless. The negative association is therefore compatible with a substitution in which an unmeasured producer increased at the expense of *B. thetaiotaomicron*.

Our study has several limitations. Because facility, cohort, and calendar time are inseparable in this design, we cannot decompose the site effect into independent mechanisms. The modest sample size (six mice per cell for the multi-omic measurements) limits the power of the interaction tests and makes individual taxon importance estimates unstable. The use of male mice only, a single arsenic dose, and a single time window for the microbiome sampling further restricts generalization, and the metabolomic associations are correlational.

Despite these limitations, this study has important implications for studies of metabolic toxicity. Standardizing exposure alone is insufficient for cross-study comparison. The environmentally driven variation in the baseline microbiome was large enough to overprint and potentially even reverse the effect of the toxicant of interest. Previous work supports the finding that toxicant-mediated outcomes are baseline-dependent and explains the inconsistency in the rodent arsenic literature, where the direction of arsenic-induced microbiome change has varied from study to study (Chi et al. 2017; Chiocchetti et al. 2019; Coryell et al. 2018). Microbial production of short-chain fatty acids, alterations in bile acids, and gut-barrier integrity all influence glucose regulation (Chi et al., 2017; Coryell et al., 2018), and each of these factors differed markedly between sites. Therefore, the baseline microbiome must be characterized, as it cannot be controlled in the absence of an experiment conducted with gnotobiotic mice.

Our findings uphold the view that the gut microbiome is an active determinant in the host response to inorganic arsenic, and that accounting for its environmentally driven variation will be essential for understanding why the same exposure can produce different metabolic outcomes. Rather than being explained by a single taxon or metabolite, our findings point to the conclusion that site-specific differences in arsenic response are driven by complementary effects of overall microbial community composition. Our results suggest that community-level characterization of microbiome composition may help to explain why arsenic-associated diabetes risk varies across human populations in exposure settings (Jorgensen et al., 2025; Spaur et al., 2024; Weiss, Sun, Jackson, Turyk, Wang, Brown, Aguilar, Hanis, et al., 2024). (Future research should be conducted to determine which subset of the gut microbiota is most important in modulating the arsenic response in order to develop interventions to mitigate the adverse effects of this widespread environmental toxicant.

## Acknowledgments

This work was supported by the National Institutes of Health (P30 ES027792, R01 ES028879, and R21 ES030884 supporting RMS and LZ). The authors would like to acknowledge Dr. Yang Chen for his assistance with microbiome data processing.

