## Supplementary Material for "Home is Where the Heterogeneity Is: Housing Facility-level Differences in the Gut Microbiome and Metabolic Phenotype Confound Arsenic Effects on Glucose Homeostasis in Male Mice"

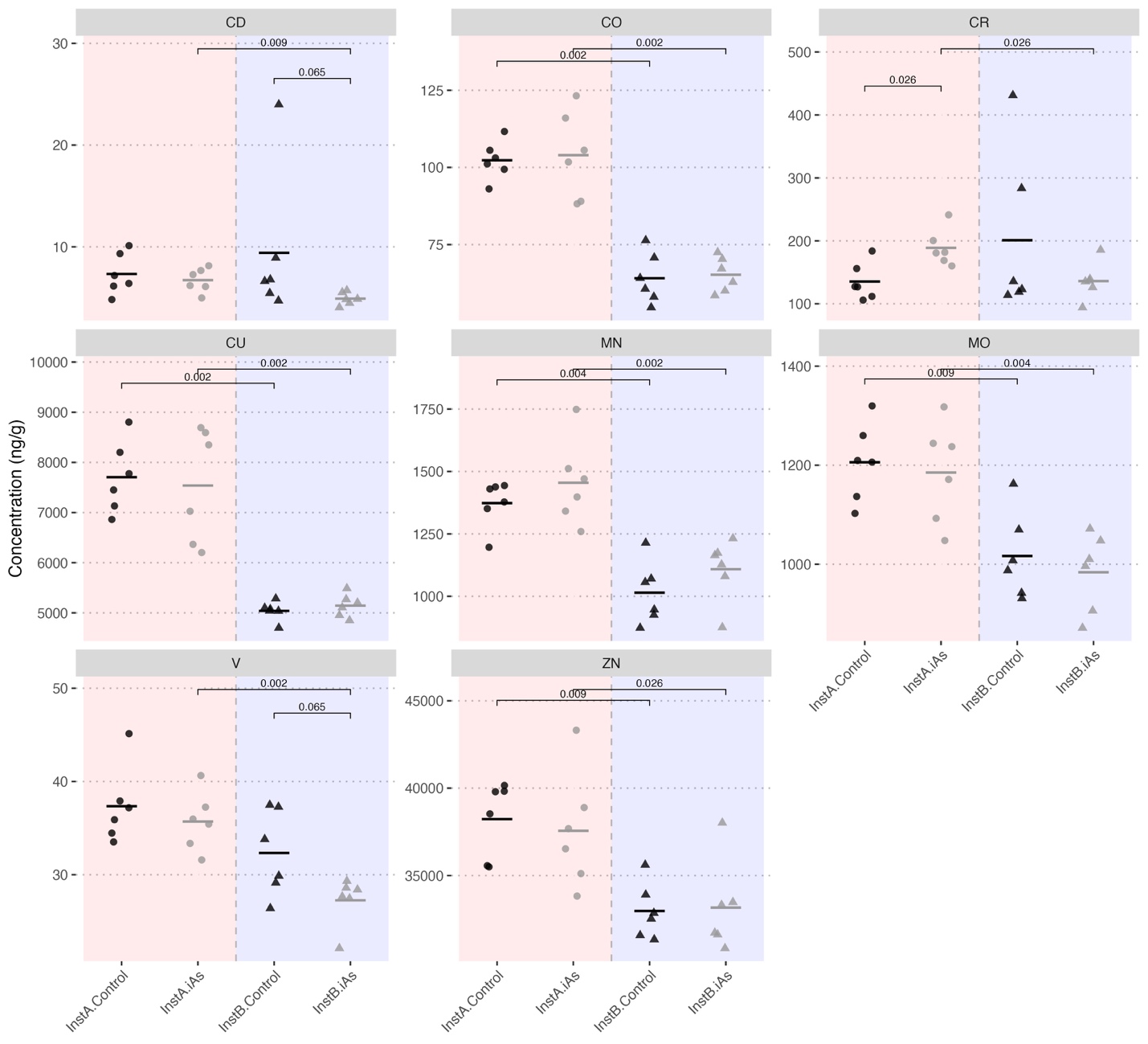

**Supplementary Figure 1*.*** *Hepatic trace metal concentrations (ng/g) measured by ICP-MS, shown by facility and treatment group. Each panel displays one element. Groups were compared by Welch's t-test; only comparisons where p < 0.1 are displayed. Horizontal crossbars indicate group means.*

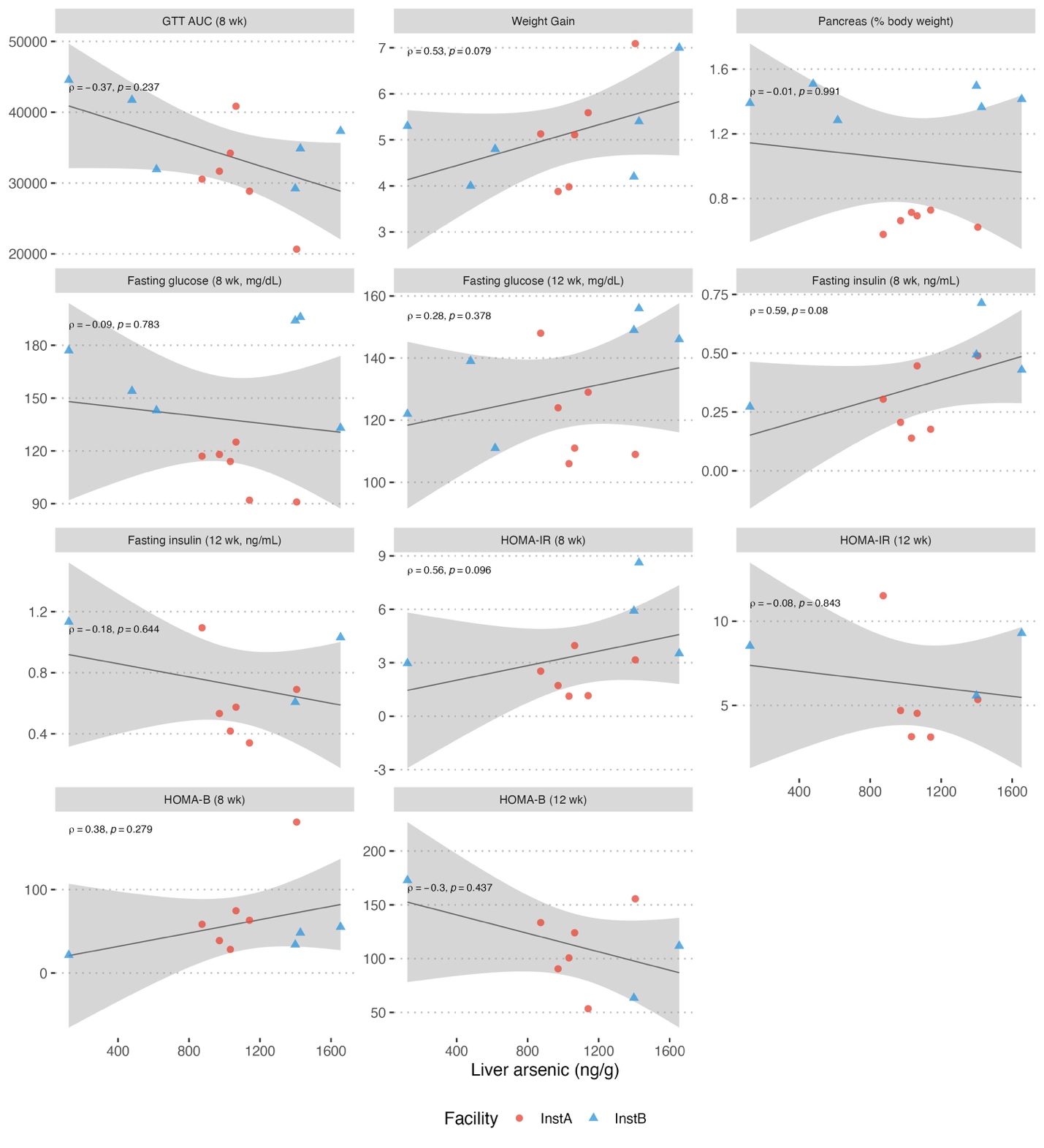

**Supplementary Figure 2*.*** *Spearman correlation between hepatic arsenic concentration (ng/g) and metabolic outcomes among iAs-treated animals only (n = 12). Panels show GTT AUC at 8 weeks, body weight gain, pancreas weight (% body weight), and fasting glucose, fasting insulin, HOMA-IR, and HOMA-β at 8 and 12 weeks. Lines are linear fits with 95% confidence bands.*

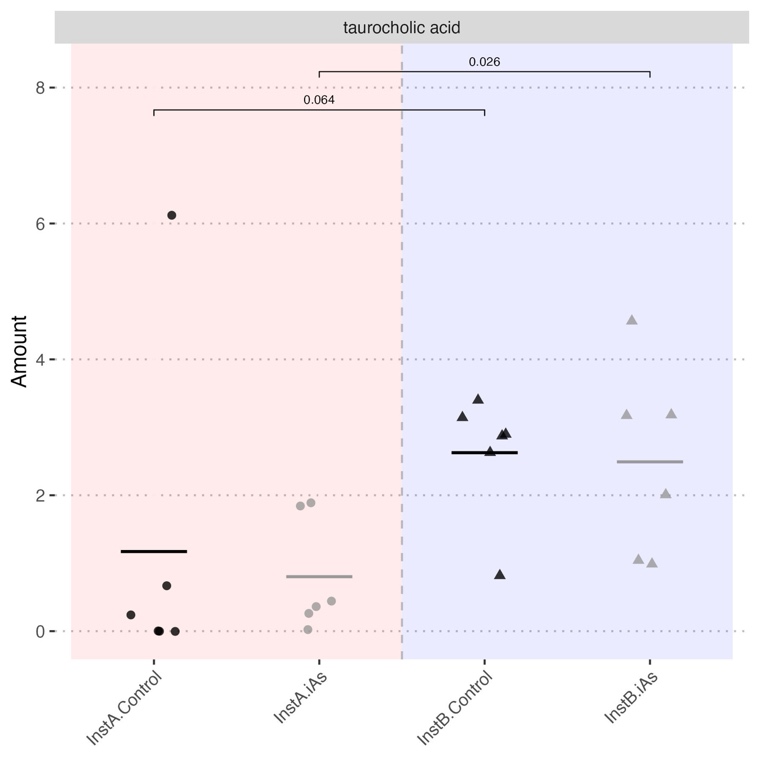

***Supplementary Figure 3.*** Targeted bile acid concentrations in cecal contents by facility and treatment group. Groups were compared by Welch's t-test; only comparisons where p < 0.1 are displayed. Horizontal crossbars indicate group means.

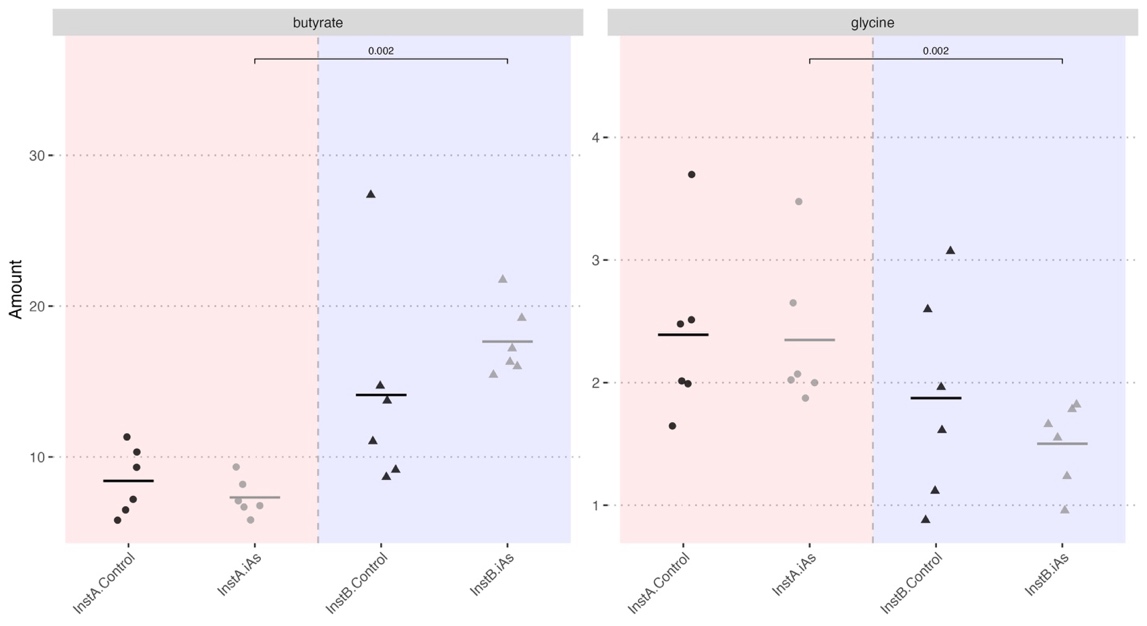

**Supplementary Figure 4.** *Targeted polar metabolite concentrations (PFBBr derivatization) in cecal contents by facility and treatment group. Groups were compared by Welch's t-test; only comparisons where p < 0.1 are displayed. Horizontal crossbars indicate group means.*

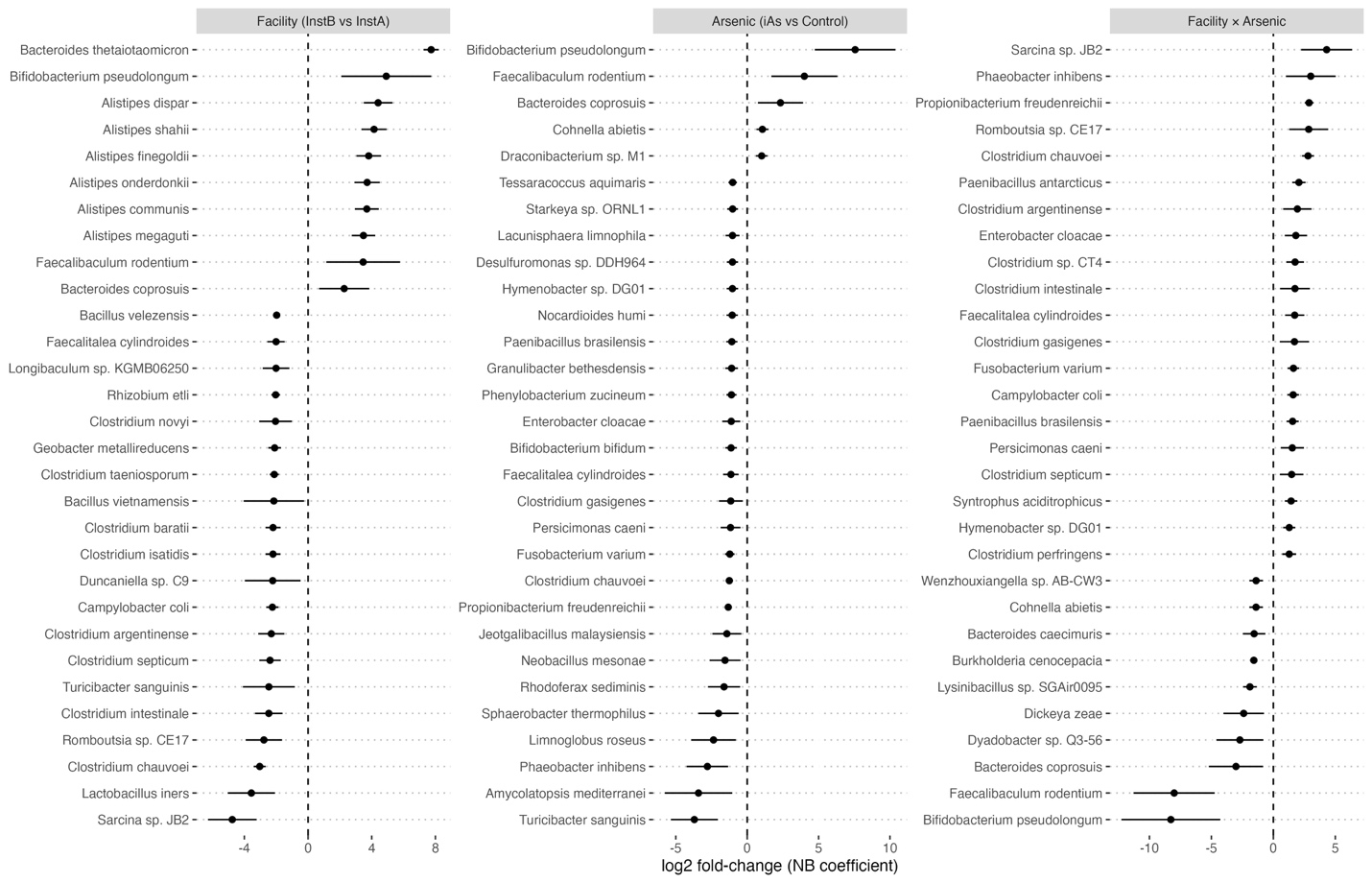

**Supplementary Figure 5*.*** *Differentially abundant taxa identified by negative binomial generalized linear models. Points show the log2 fold-change for each significant taxon with 95% confidence intervals, grouped by model term (facility, treatment, and facility × treatment interaction). Taxa were retained at adjusted p < 0.05.*

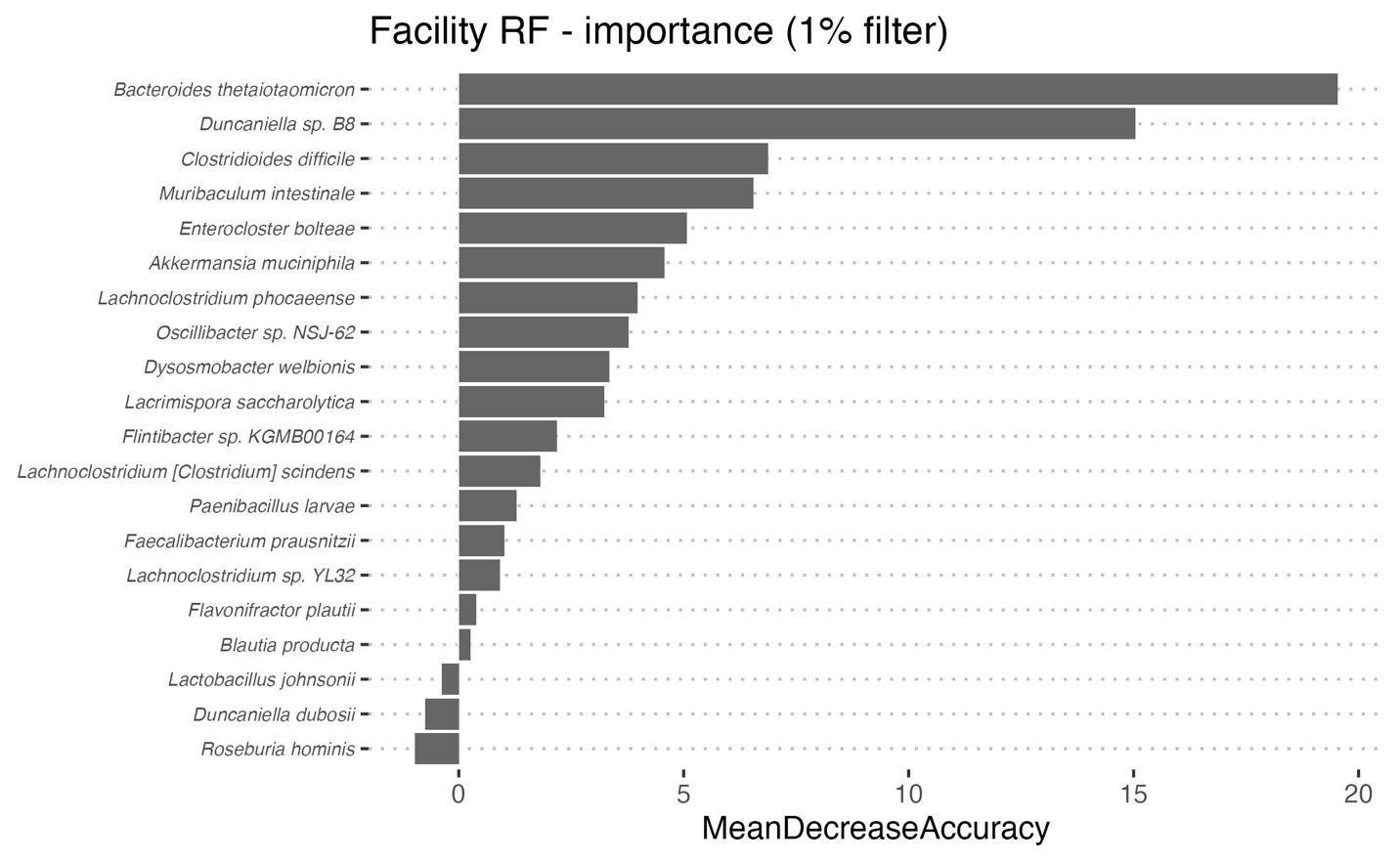

**Supplementary Figure 6.** *Variable importance for the random forest facility classifier at the 1% mean relative abundance filter (22 taxa). Bars show mean decrease in accuracy from the full-data fit. Model performance across all abundance filters is reported in Table 2.*

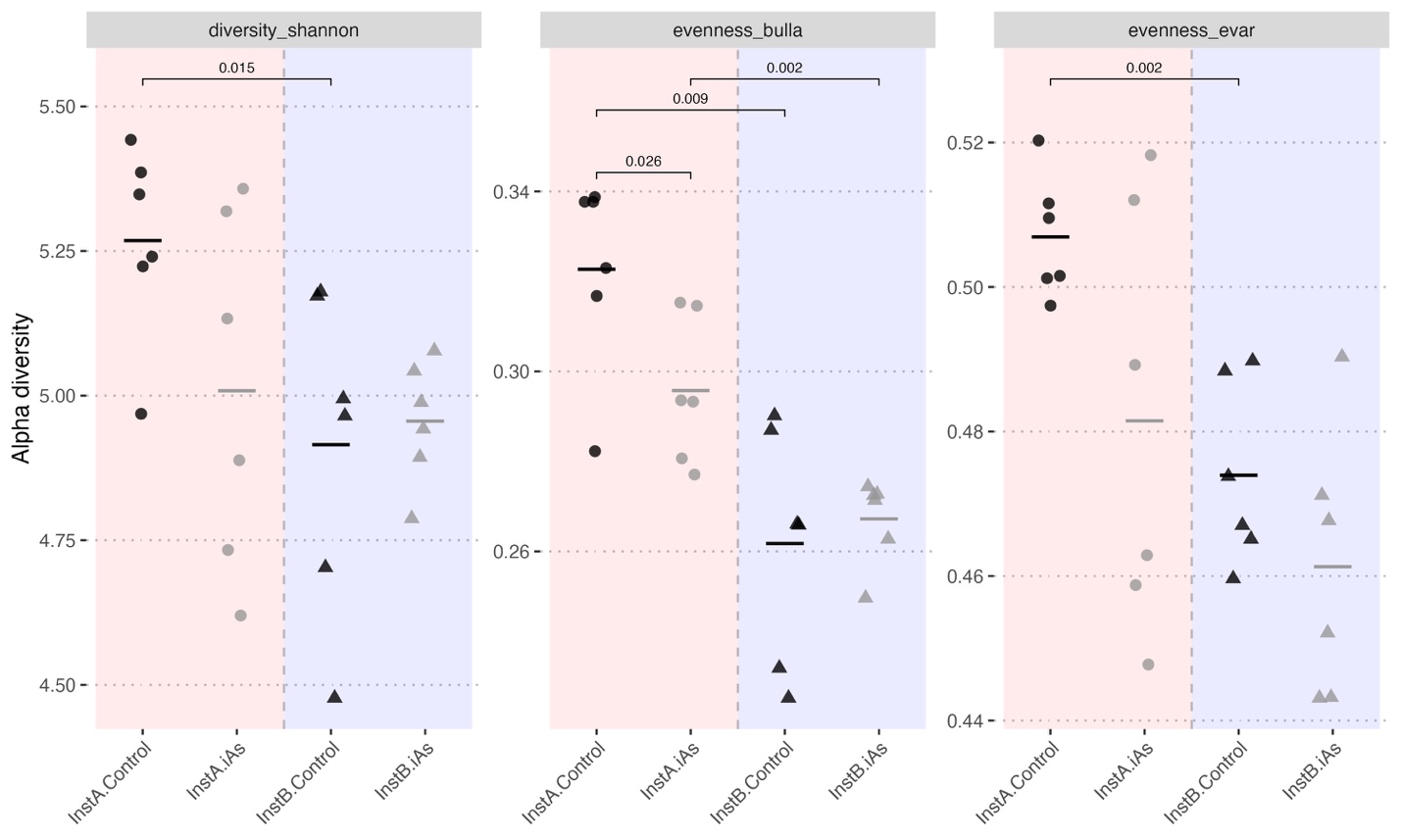

**Supplementary Figure 7.** Alpha diversity metrics that differed significantly between groups: Shannon diversity, Bulla evenness, and eVar evenness. Indices were computed on rarefied data to account for differences in sequencing depth between facilities. Groups were compared by Wilcoxon rank-sum test; only comparisons where p < 0.1 are displayed. Horizontal crossbars indicate group means.

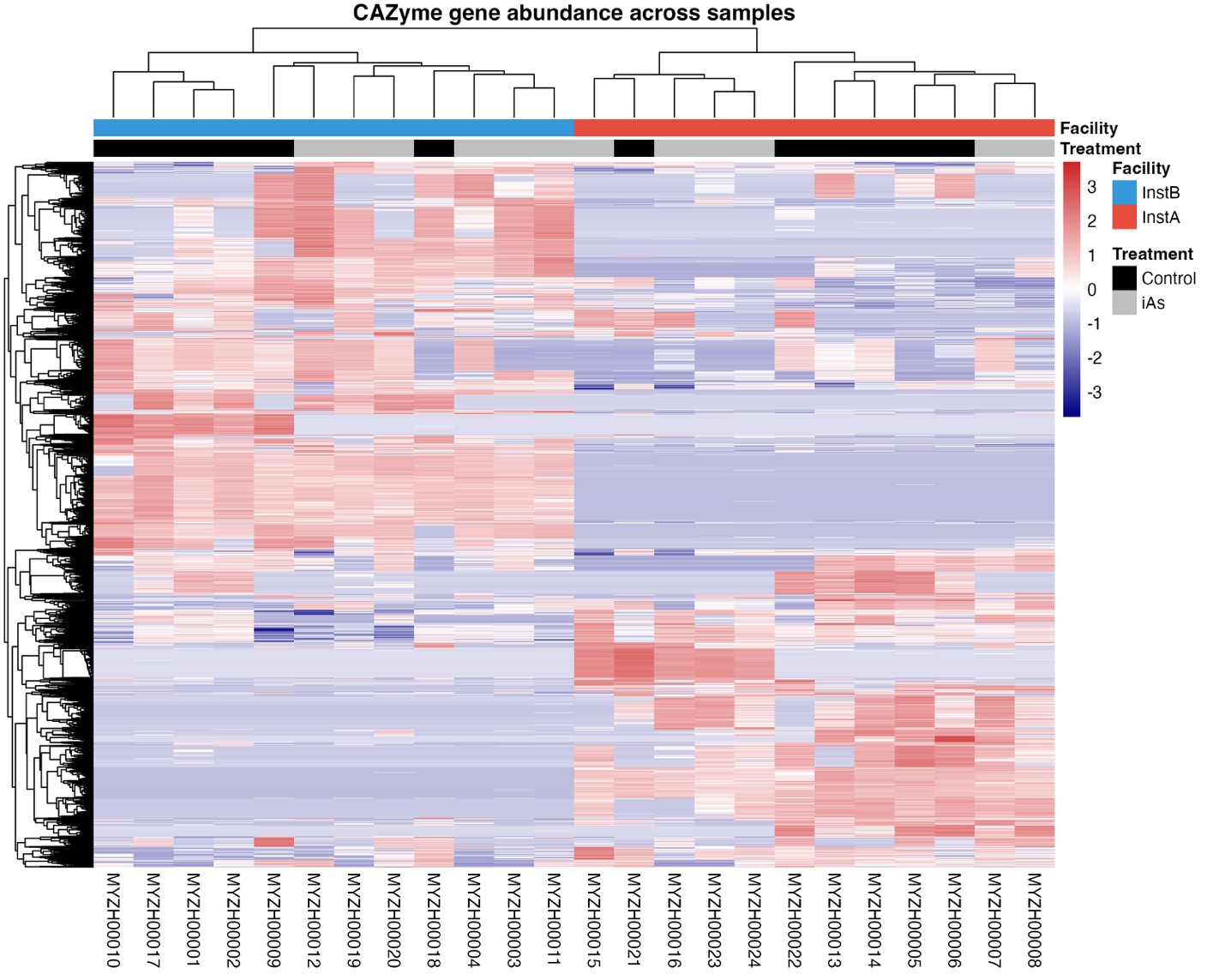

**Supplementary Figure 8.** *Hierarchically clustered heatmap of carbohydrate-active enzyme (CAZyme) gene abundances across samples, annotated with run_dbcan. Cell color encodes the row-wise z-score of normalized gene counts. Column annotations indicate facility and treatment.*

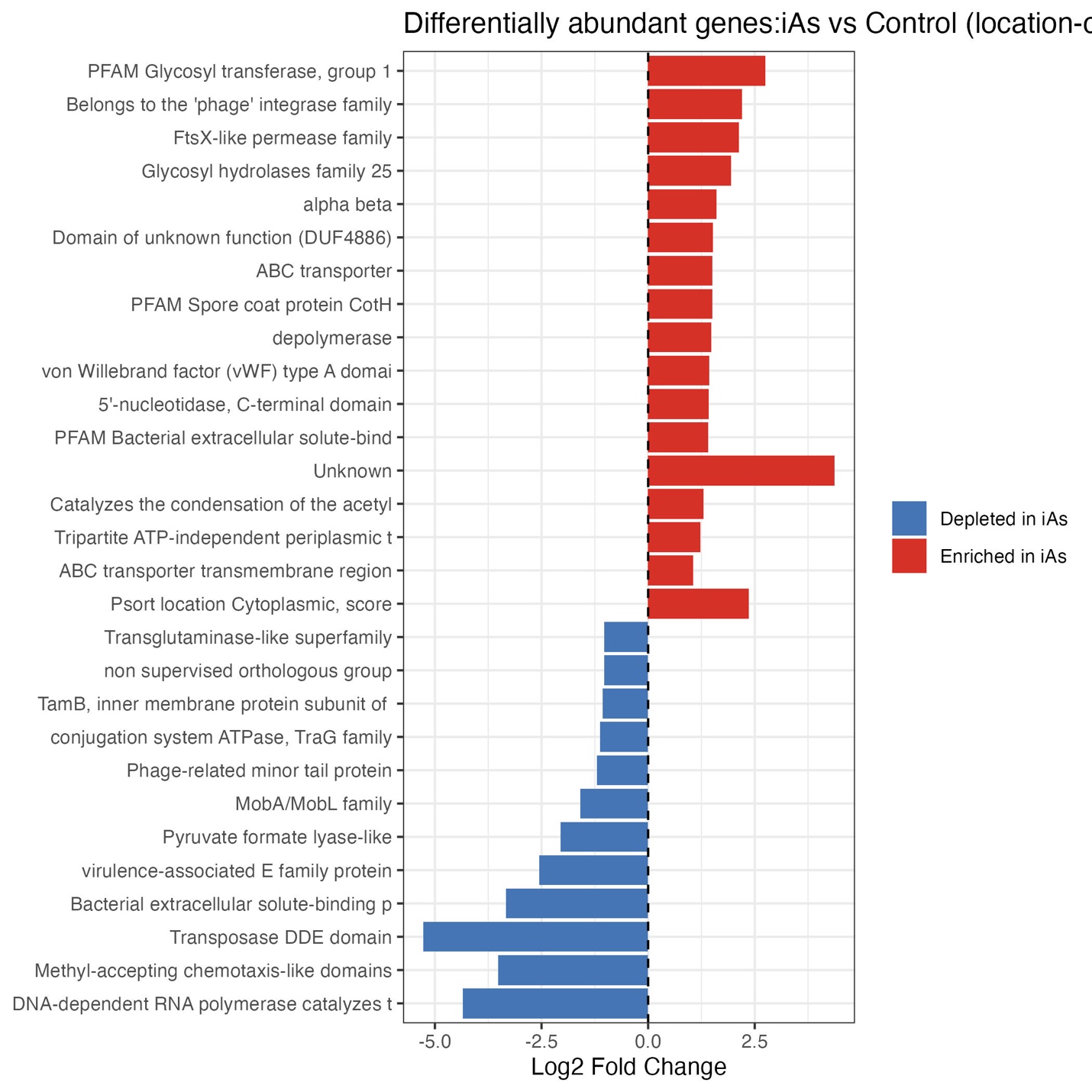

**Supplementary Figure 9.** *Genes differentially abundant between iAs-treated and control animals, controlling for facility (BH-adjusted p < 0.05, |log2 fold-change| > 1), labeled by functional annotation. Bars are colored by direction of effect (red, enriched under iAs; blue, depleted under iAs). None of the 139 differentially abundant genes were annotated as arsenic-metabolism genes in AsgeneDB.*

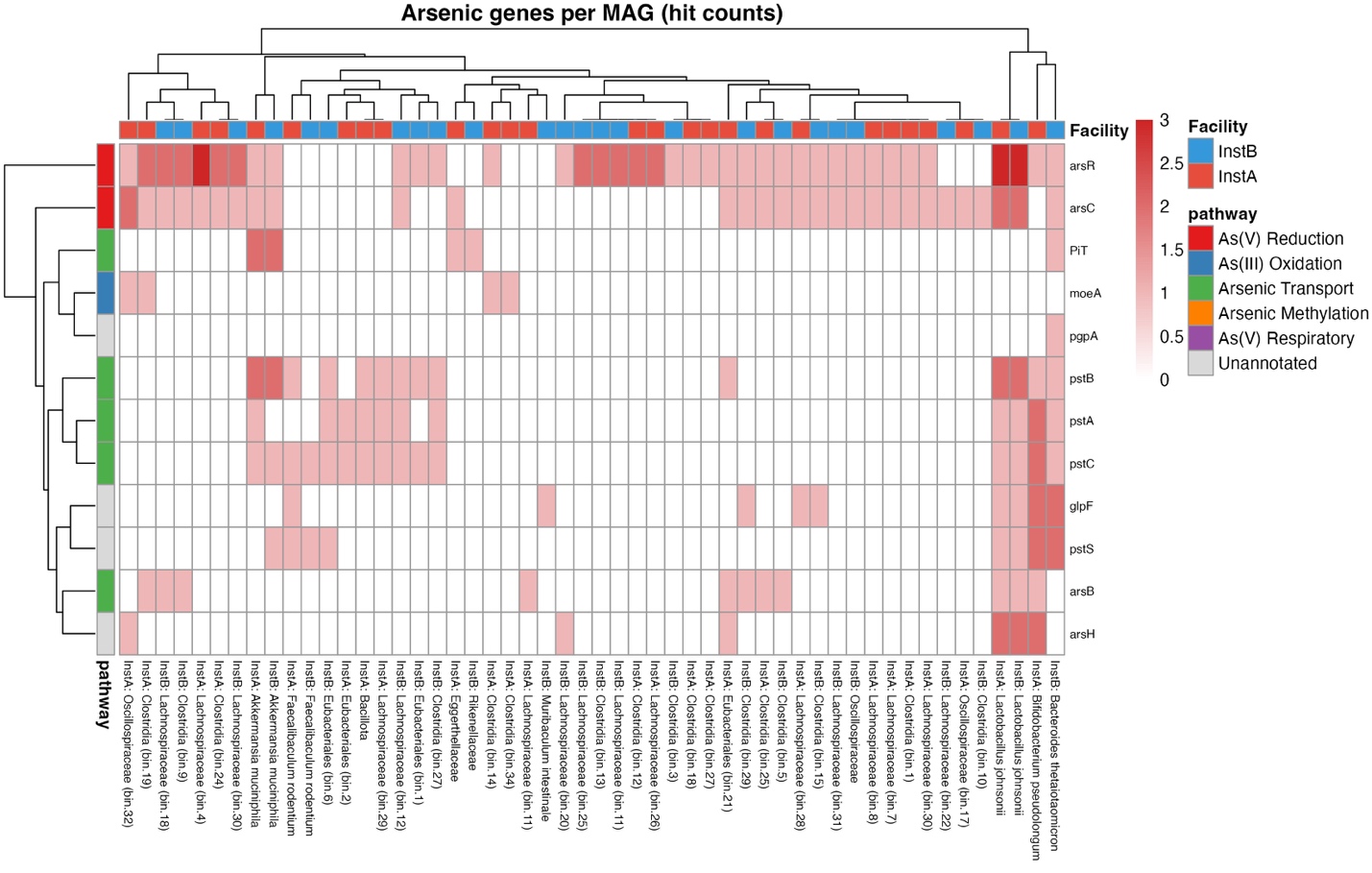

**Supplementary Figure 10.** *Arsenic-metabolism gene content of metagenome-assembled genomes (MAGs), assembled separately at each facility. Cell color encodes the number of AsgeneDB hits per gene family within each MAG. Row annotation indicates the arsenic-cycling pathway assigned to each gene family; column annotation indicates the facility of assembly. MAGs are labeled by facility and the lowest confident taxonomic assignment.*
